# High-Resolution Structure of the Nuclease Domain of the Human Parvovirus B19 Main Replication Protein NS1

**DOI:** 10.1101/2021.12.20.473600

**Authors:** Jonathan L. Sanchez, Niloofar Ghadirian, Nancy C. Horton

## Abstract

Two new structures of the N-terminal domain of the main replication protein, NS1, of Human Parvovirus B19 (B19V) are presented. This domain (NS1-nuc) plays an important role in the “rolling hairpin” replication of the single-stranded B19V DNA genome, recognizing origin of replication sequences in double-stranded DNA, and cleaving (*i.e*. nicking) single-stranded DNA at a nearby site known as the trs. One structure of NS1-nuc is solved to 2.4 Å and shows the positions of two bound phosphate ions. A second structure shows the position of a single divalent cation in the DNA nicking active site. The threedimensional structure of NS1-nuc is well conserved between the two forms, as well as with a previously solved structure of a sequence variant of the same domain, however shown here at significantly higher resolution. Using structures of NS1-nuc homologues bound to single- and double-stranded DNA, models for DNA recognition and nicking by B19V NS1-nuc are presented which predict residues important for DNA cleavage and for sequence specific recognition at the viral origin of replication.

## Introduction

Human Parvovirus B19 (B19V) is a ubiquitous virus infecting the majority of the human population^*1, 2*^. B19V, a *Parvoviridae* family member of the genus *Erythrovirus* has been associated with a myriad of different illnesses; B19V was first discovered as the cause of aplastic crisis in patients with chronic hemolytic anemia^*3*^, then as the causative agent of *erythema infectiosum* (Fifth disease) in 1983^*4*^ which results in mild fever and a distinctive rash in children, and fever often with hepatitis and arthralgia in adults. B19V infection is also associated with pure red-cell aplasia from persistent infection in immunocompromised patients, and *hydrops fetalis* (a serious condition of the fetus) in pregnant women^*1*^. In addition, B19V infection has also been associated with other serious conditions such as inflammatory cardiomyopathy and the induction of autoimmune or autoimmune-like disease (short or long term)^*5–8*^.

B19V is a single-stranded nonenveloped DNA virus of 5596 nucleotides, with an internal coding region flanked by palindromic sequences capable of forming terminal hairpin structures. B19V replicates in erythroid progenitors using a rolling hairpin mechanism^*9*^. The viral genome encodes six protein products: VP1 and VP2 which compose the viral capsid, NS1, the main replication protein, and three smaller nonstructural proteins^*10–12*^. Viral replication utilizes cellular factors, proposed to be coordinated by the viral NS1 protein^*13–15*^, and is thought to make use of the terminal hairpins^*13, 15*^ **(Fig. 1A**). Extension of the 3’ end of one terminal hairpin by a cellular polymerase results in replication of the majority of the viral genome (Step 1, **Fig. 1A**), and NS1 is thought to bind to repeat sequences (NSBE1-4)(**Fig. 1B**) and nick or cleave in one strand at the trs (the “terminal resolution site”)(Step 2, **Fig. 1A**), producing a new 3’ end that can be used to prime synthesis of the remaining viral DNA (Step 3, **Fig. 1A**)^*14*^. Based on amino acid sequence and homology to other parvoviral replication proteins, B19V NS1 (NS1) is predicted to contain both nuclease and helicase domains involved in B19V replication. NS1 also contains a C-terminal domain involved in the protein’s promoter transactivation activity^*16*^ which acts upon its own viral promoter, p6, as well as several host promoters^*17–21*^. Furthermore, the genome of B19V is known to insert into host DNA, a reaction likely involving double-stranded DNA (dsDNA) recognition and strand nicking by NS1 followed by host DNA repair^*22*^.

**Figure 1.**
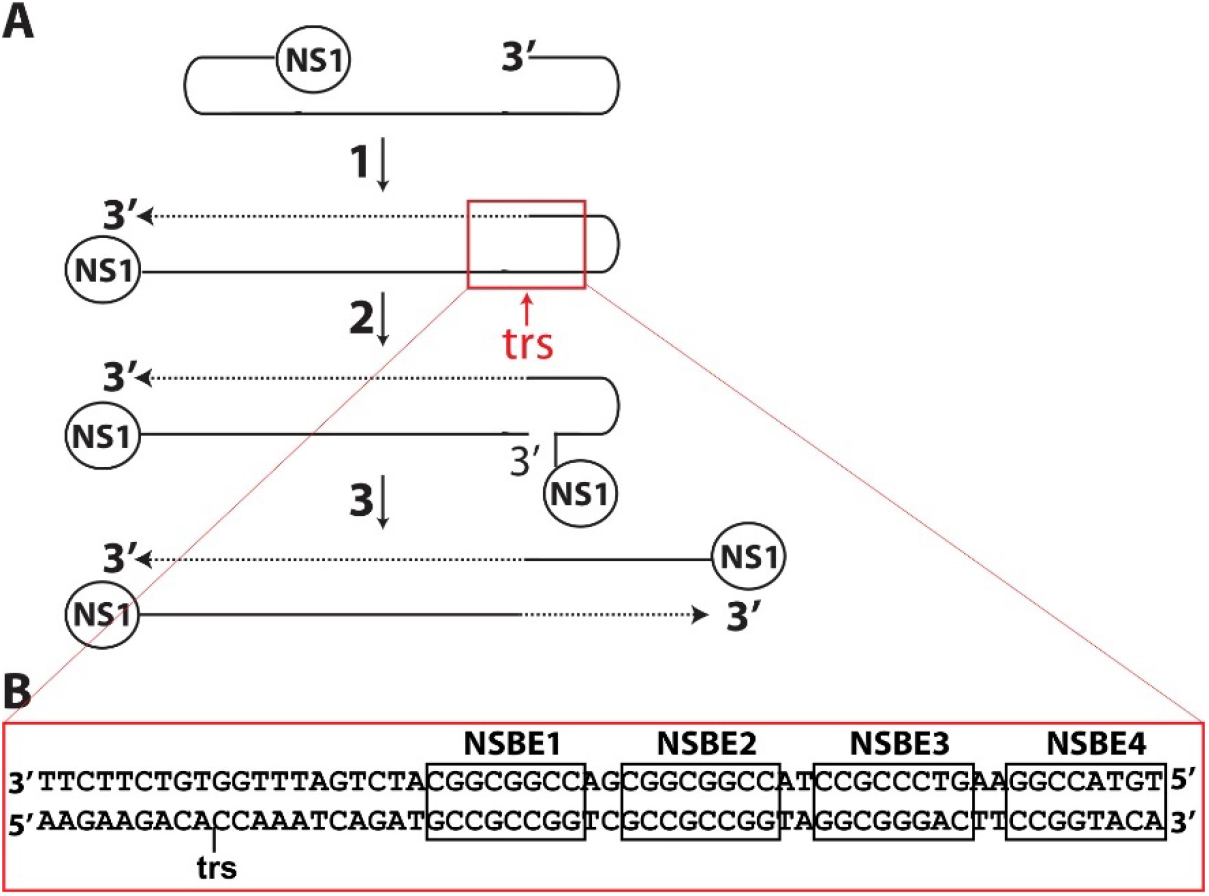
A rolling hairpin model of viral genome replication and viral origin of replication sequences. **A.** Replication using the cellular machinery is primed by the 3’ folded over hairpin (or ITR). Upon double-stranded formation, transcription proceeds producing new copies of NS1. NS1 binds at the GC rich sequences denoted NSBE1-4 and cleaves at the nearby terminal resolution site or trs, leaving NS1 covalently attached to the 5’ end at the site of cleavage. Replication of the remaining segment of the genome is completed using the newly generated 3’OH and following unfolding of the terminal hairpin. **B.** DNA sequences at the viral origin of replication including the trs and NSBE sequences.

Prior studies show that the isolated N-terminal nuclease domain of B19V NS1 (NS1-nuc) binds sequence specifically to the NSBE sequences in dsDNA, and is also responsible for sequence specific DNA cleavage (or nicking) in single-, but not double-stranded DNA, at the trs site^*23*^. Since DNA encountered by NS1 is doublestranded, it is assumed that binding of NS1 to its target sequences in dsDNA induces strand separation nearby, allowing for the endonuclease activity of NS1 to cleave at the trs site. However, how NS1 recognizes its target sequences in dsDNA as well as at the nicking site are currently unknown, as is the mechanism by which NS1 binding to DNA at the NSBE induces strand separation. To begin to answer these questions, we determined the high resolution crystal structure of NS1-nuc. Two crystal forms were solved, one at 2.4 Å resolution (Form I) which shows binding to phosphate ions at the predicted double- and single-stranded DNA binding sites, and another (Form II) at 3.5 Å resolution which shows binding of a single Mg^2+^ in the DNA nicking active site. Using homology modeling, three dimensional models for 1) trs sequence recognition in single-stranded DNA (ssDNA), 2) DNA cleavage in ssDNA, and 3) dsDNA binding at the NSBE sequences are presented.

## Materials and Methods

### Protein expression and purification

Purification of Form I NS1-nuc (residues 2-176 of B19V NS1, see Supplemental Information) free of purification tags was performed as previously described^*23*^ using an N-terminal 6xhistidine and MBP-tagged fusion protein with a TEV protease cleavage site between the tags and the NS1-nuc sequences (see Supplementary Information). Following cell lysis using an Avestin Emulsiflex C3, cell debris were pelleted and the cell-free lysate incubated with pre-equilibrated Talon resin (Clonetech, Inc.). The partially pure eluted protein was then incubated with a 1:1 molar ratio of TEV protease^*24*^ overnight at 4 °C. NS1-nuc free of MBP- and his-tags was then further purified with DEAE and Heparin FPLC (GE, Inc.). A longer construct of NS1-nuc (residues 2-209 of B19V NS1, see Supplementary Information) was used in the Form II crystals and was prepared with an N-terminal 6xhistidine tag and expressed in Tuner (DE3) cells overnight at 17°C following induction. Cells were lysed using an Avestin Emulisflex C3, then centrifuged to pellet cell debris. Purification proceeded with Talon resin (Clonetech, Inc.) chromatography followed by DEAE FPLC (GE). Purified protein was dialyzed into 0.1 M bis-tris-propane pH 9.5, 150 mM NaCl, 1 mM 2-mercaptoethanol, and 50% glycerol, aliquoted, flash frozen in liquid nitrogen, then stored at −80°C until needed.

### Crystallization, x-ray diffraction data collection, structure solution, and refinement

Crystallization proceeded using the hanging drop vapor diffusion method. NS1-nuc protein was dialyzed extensively against 0.1 M bis-tris-propane pH 9.5, 150 mM NaCl, 1 mM 2-mercaptoethanol and concentrated to 5-10 mg/ml. Form II crystals appeared with the crystallization solution consisting of 2.5 M NaCl, 0.1 M Tris-HCl pH 7.0, and 200 mM MgCl2 after several weeks at 4°C. For x-ray diffraction data collection, crystals were harvested and exchanged into 2.5 M NaCl, 0.1 M Tris-HCl pH 7.0, and 30% glycerol then flash frozen and stored in liquid nitrogen. Form I crystals appeared after several weeks at 17°C in the crystallization solution containing 15% PEG 3350, 0.1 M sodium citrate pH 4.5, 0.1 M NaCl, and 0.1 M LiCl and a 7:1 molar ratio of NS1-nuc to NSBE containing DNA (see Supplementary Information, this dsDNA contains two NS1 binding element repeats found at the B19V origin of replication). Form II NS1-nuc crystals were exchanged into cryoprotectant (15% PEG 3350, 0.1 M sodium citrate pH 4.5, 0.1 M NaCl, 0.1 M LiCl, and 30% glycerol) prior to flash freezing in liquid nitrogen. X-ray diffraction data collection was performed at SSRL BL 9-2 at 100K using Blu-Ice software^*25*^. Data processing, including integration, scaling, and merging, was performed with iMosflm^*26*^ and SCALA^*27, 28*^. Structure solution of Form I NS1-nuc was performed using molecular replacement in PHASER^*29*^ within the PHENIX software suite^*30*^, and by searching for one copy of the nuclease domain using coordinates from PDB accession code 6USM^*31*^. Structure building and refinement proceeded through an iterative process using COOT^*32, 33*^ and refinement using PHENIX^*30, 34–36*^. Solution of Form II proceeding using the same procedure, however with the Form I structural coordinates as the search model. Refinement of Form II made use of Form I as a reference structure, as well as secondary structure geometry restraints. RMSD calculations were performed with UCSF Chimera^*37*^, Pymol (Schrodinger), and DALI^*38*^. Images of structural models and electron density were prepared with UCSF Chimera^*37*^ and Pymol (Schrodinger). Electrostatic calculations performed with the software APBS^*39*^ in Pymol (Schrodinger). Topology diagrams were made with PDBsum^*40*^.

### Sequence variation analysis of NS1-nuc

Sequences (195 total) of Human Parvovirus B19 NS1 (residues 1-176) were extracted from NCBI using BLASTp and the sequence of Form I NS1-nuc as the search sequence. WebLOGO^*41*^ was used to create a figure to display amino acid sequence variations.

## Results and Discussion

### Structure Solution, Refinement, and Overall Analysis

Crystallographic data and structure refinement statistics are shown in **Table 1**. X-ray diffraction data were collected, scaled, and truncated to resolutions based on scaling, signal to noise, cross-correlation analysis, and final structure and map quality after structure refinement. These resolutions are 2.4 Å for Form I, where the highest resolution shell shows I/σ of 2.4, CC_1/2_ of 38.5%, and final R_work_ and R_free_ of 20.0% and 25.5%, respectively. In the case of Form II, data were truncated at 3.5 Å where the highest resolution shell shows I/σ of 1.67, CC1/2 of 89%, and final R_work_ and R_free_ of 26.1% and 29.6%, respectively. Electron density for DNA was not evident in the Form I maps, despite its addition to the protein solution used in the crystallization experiments. The additional residues on the C-terminus of the NS1-nuc construct used for Form II crystallization were also not evident in the electron density maps, possibly due to protein degradation during crystallization (**Fig. S1**). **Figure 2A** shows two views of the NS1-nuc ribbon diagram of Form I, and **Figures 2B-C** show secondary structural elements and other notable structural features. The largest differences between Forms I and II NS1-nuc structures include a variation in the trace at residues 24-30 (**Fig. S2A-C**), as well as the absence of residues 127-128 in Form II (**Fig. S2A**), which are residues implicated in dsDNA binding (discussed further below). These structural differences may originate from crystal packing interactions. One phosphate ion per asymmetric unit was found in the Form I structure (orange spheres, **Fig. 2A**, orange and yellow stick, **Fig. 3A-B**) which interacts with residues of two neighboring NS1-nuc copies in the crystal lattice (**Fig 3B, Fig. S3**). The phosphate ion is within salt bridging distance to His81 (3.8 Å) of one NS1-nuc copy (red, **Fig. 3B**) and Lys119 (3.4 Å) of another (Lys119’, **Fig. 3B**). The phosphate ion also interacts through water molecules to three side chains: Tyr141 and His83 of one copy of NS1-nuc, and Try130 of the other (Tyr130’, **Fig. 3B**). In the Form II crystal structure, significant (*i.e*. 3σ) positive difference density was seen in the position occupied by Zn^2+^ in a previously determined structure of NS1-nuc (PDB accession code 6USM^*31*^) suggesting divalent cation binding (2Fo-Fc at 1σ map in blue, 1Fo-Fc map in magenta at 3σ when Mg^2+^ is omitted from the model, **Fig. 3C**). Mg^2+^ was modeled into this position (green sphere, **Fig. 3A**) since crystallization conditions contained 200 mM Mg^2+^.

**Figure 2.**
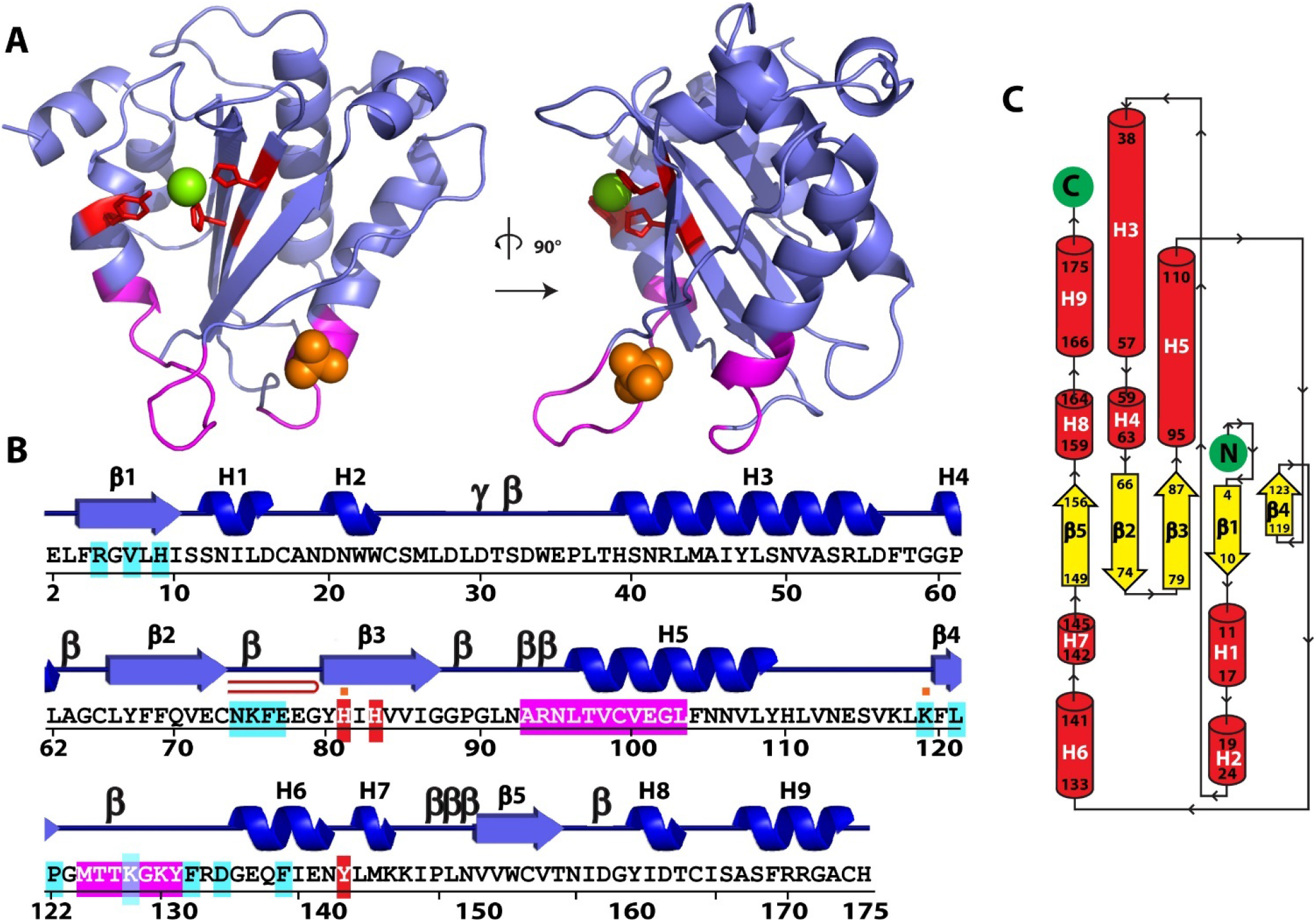
Overall Structure of Human Parvovirus B19 NS1 nuclease domain (NS1-nuc). **A.** Overview of Form I NS1-nuc with the positions of putative DNA cleavage active site residues in red and predicted dsDNA binding residues shown in magenta. Mg^2+^ identified in Form II shown as green sphere, and a phosphate ion bound to Form I shown as orange spheres. **B.** Sequence, secondary structure, and notable features of Form I NS1-nuc. H1-H9: alpha helices, β1-β5: β-strands, β: β-turn, γ: γ-turn, red hairpin: β-hairpin, white letters in red boxes: putative DNA cleavage active site residues (*i.e*. active site residues), white in magenta boxes: predicted dsDNA binding residues, cyan boxes: predicted ssDNA binding residues, small orange boxes above sequence: residues in contact with bound phosphate ions. **C.** Topology diagram of Form I NS1-nuc with α helices as red cylinders, β strands as white arrows, N and C termini in green circles, and residue number of secondary structural elements in black.

**Figure 3.**
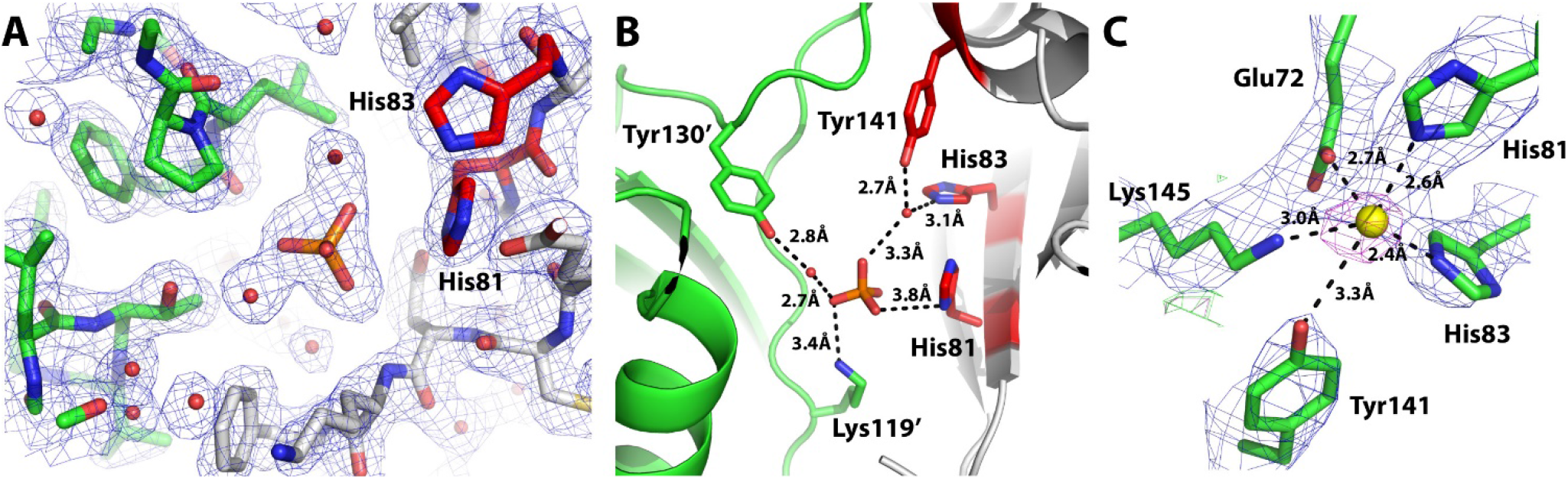
Ligands found bound to NS1-nuc structures. **A.** A phosphate ion is bound between two neighboring copies of NS1-nuc (shown in green/blue/red and white/blue/red with the two active site residues of the white NS1-nuc copy shown in red/blue). 2Fo-Fc map shown at 1σ in blue. **B.** As in A, with hydrogen bonding and salt bridging interactions between the bound phosphate ion and residues from the two different copies of NS1-nuc **C.** A 3σ peak is found in the Form II 1Fo-Fc map (magenta, modeled atom as yellow sphere) near five residues of the active site and modeled as Mg^2+^ (yellow sphere). 2Fo-Fc map shown at 1σ in blue.

**Table 1.** Data collection and refinement statistics.

|  | **Form I** | **Form II** |
| --- | --- | --- |
| **PDB accession code** | 7SZY | 7SZX |
| **Site of data collection** | SSRL BL9-2 | SSRL BL9-2 |
| **Wavelength** | 0.97946 Å | 0.97946 Å |
| **Resolution range** | 45.06 Å - 2.4 Å (2.486 Å - 2.4 Å) | 46.21 Å - 3.5 Å (3.625 Å - 3.5 Å) |
| **Space group** | P 2 21 21 | P 32 2 1 |
| **Unit cell** | 45.0599 Å, 49.0999 Å, 74.9899 Å | 106.71 Å, 106.71 Å, 59.5904 Å |
|  | 90° 90° 90° | 90° 90° 120° |
| **Total reflections** | 13,532 (1322) | 10,152 (976) |
| **Unique reflections** | 6,799 (665) | 5,099 (488) |
| **Multiplicity** | 2.0 (2.0) | 2.0 (2.0) |
| **Completeness (%)** | 98.44 (98.66) | 98.70 (97.41) |
| **Mean I/sigma(I)** | 7.21 (2.43) | 5.37 (1.67) |
| **Wilson B-factor** | 28.31 | 91.66 |
| **R-merge** | 0.1022 (0.5331) | 0.07577 (0.4226) |
| **R-meas** | 0.1446 (0.7539) | 0.1072 (0.5977) |
| **CC<sub>1/2</sub>** | 0.981 (0.385) | 0.993 (0.89) |
| **CC\*** | 0.995 (0.745) | 0.998 (0.971) |
| **Reflections used in refinement** | 6,797 (665) | 5,074 (488) |
| **Reflections used for R-free** | 333 (40) | 508 (50) |
| **R-work** | 0.2001(0.2677) | 0.2605 (0.3421) |
| **R-free** | 0.2548 (0.2810) | 0.2956 (0.4067) |
| **CC(work)** | 0.951 (0.772) | 0.908 (0.759) |
| **CC(free)** | 0.907 (0.631) | 0.825 (0.685) |
| **Number of non-hydrogen atoms** | 1432 | 1294 |
| **macromolecules** | 1337 | 1289 |
| **ligands** | 5 | 1 |
| **solvent** | 90 | 4 |
| **Protein residues** | 174 | 171 |
| **RMS(bonds)** | 0.002 | 0.004 |
| **RMS(angles)** | 0.52 | 0.87 |
| **Ramachandran favored (%)** | 95.93 | 94.01 |
| **Ramachandran allowed (%)** | 4.07 | 5.39 |
| **Ramachandran outliers (%)** | 0.00 | 0.60 |
| **Rotamer outliers (%)** | 1.43 | 0.00 |
| **Clashscore** | 2.29 | 6.09 |
| **Average B-factor** | 29.13 | 94.33 |
| **macromolecules** | 28.94 | 94.51 |
| **ligands** | 41.05 | 90.01 |
| **solvent** | 31.26 | 37.16 |

Statistics for the highest-resolution shell are shown in parentheses.

### Comparison to other parvoviral Rep protein nuclease domains

Form I NS1-nuc was solved by molecular replacement using the NS1-nuc domain found in the PDB file with accession code 6USM^*31*^. After refinement these two structures of NS1-nuc were found to be very similar, with an RMSD of 0.8 Å over 164 residue C_α_ atoms (**Fig. S4A)**, and the largest differences in structure were found in loop segments at the exterior of the protein (thick red ribbons, **Fig. S4B**). The loop segment containing residues 147-148 is not present in the 6USM coordinates, but is well ordered in Form I NS1-nuc and contains a cis-proline (**Fig. S4C**). In addition, residues at the C-terminus of the domain are truncated to 171 in 6USM, but extend to 175 in Form I NS1-nuc. A single Zn^2+^ is located in the active site of 6USM in a nearly identical position as the Mg^2+^ ion modeled in Form II NS1-nuc (**Fig. S4D**). Differences in the amino acid sequences of NS1-nuc construct used in our studies and that of 6USM occur in thirteen positions (red text, **Fig. S4E**). These originate from differences in biologically relevant sequences present in the NCBI data base (6USM follows the sequence of a laboratory isolate, Genbank entry AAG00943, Form I and II NS1-nuc follow the sequence of an isolate from a blood bank, Genbank entry ABN45789.1), and represent the two major variants present in NCBI (**Fig. S5**). These substitutions are largely conservative in nature, and no large perturbations to the two structures are found at these positions (**Fig. S6**).

**Figure 4** compares the structure of NS1-nuc (Form I, blue, **Fig. 4A-C**) to homologous parvoviral structures present in the PDB originating from Adeno Associated Virus (AAV, PDB accession code 5DCX^*42*^)(magenta, **Fig. 4A**), Minute Virus of Mice (MVM, PDB accession code 3WRN^*43*^)(yellow, **Fig. 4B**), and Human Bocovirus (HBov, PDB accession code 4KW3^*44*^)(green, **Fig. 4C**). Ribbon diagrams of pairwise superpositions (using C_α_ atoms) are shown in **Figures 4A-C**, with root mean square deviation values (RMSD using C_α_ atoms) mapped onto the structure of NS1-nuc (Form I) in **Figures 4D-F** (thicker, redder lines indicate greater RMSD, see legend in Å below each ribbon diagram). Most differences in structure occur to the positioning of loops and α-helices behind (as shown in the figure) the central β-sheet containing the active site histidine residues (marked in red in **Fig. 4A-C**). The structure of the AAV homolog contains an additional segment corresponding to the linker residues located between the N-terminal nuclease domain and the central helicase domain of AAV Rep (the homolog of B19V NS1), which are not present due to sequence truncation in the other domain structures (arrow, **Fig. 4A**). The overall RMSD values between NS1-nuc (Form I) and the other three structures are very similar, with 3.2 Å, 3.2 Å, and 3.0 Å over 163-165 residue B19V NS1-nuc C_α_ atoms and the nuclease domains of AAV Rep, MVM NS1, and HBov NS1, respectively.

**Figure 4.**
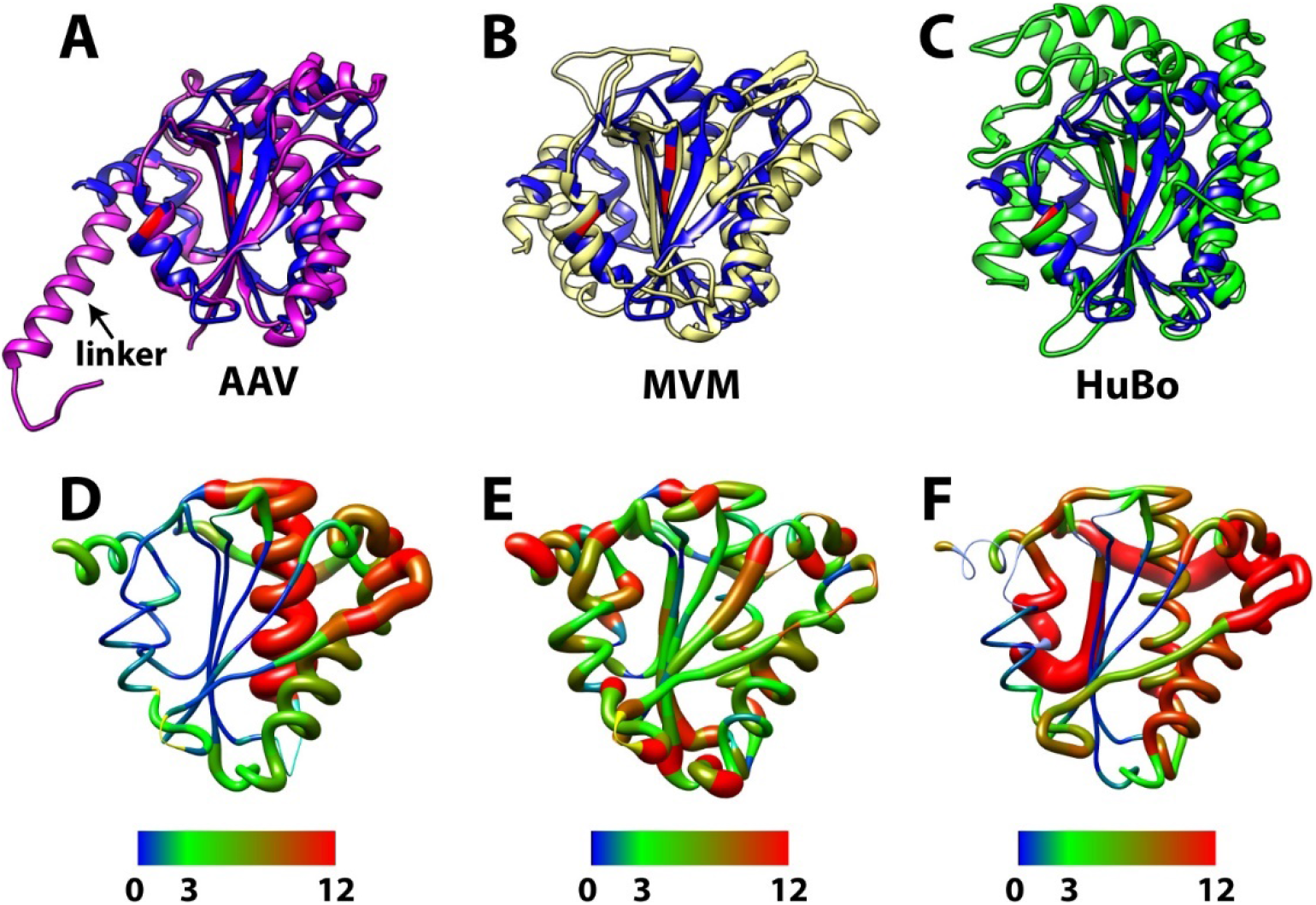
Comparison of NS1-nuc with parvoviral homologues. **A.** Comparison of Form I NS1-nuc (blue) with the nuclease domain of Adeno Associated Virus (AAV) Rep (magenta, PDB accession code 5DCX^*42*^). Positions of active site residues in NS1-nuc (Tyr141, His81, His83) shown in red. **B.** As in A, but comparing NS1-nuc (blue) to the nuclease domain of Minute Virus of Mice (MVM) NS1 (yellow, PDB accession code 3WRN^*43*^). **C.** As in A, but comparing NS1-nuc (blue) the nuclease domain of Human Bocavirus (HBov) NS1 (green, PDB accession code 4KW3^*44*^). **D.** Ribbon diagram showing RMSD between C_α_ of NS1-nuc and AAV Rep-nuc. Ribbon color and thickness indicate local RMSD, scale show below, in Å. **E.** As in D but NS1-nuc and MVM NS1-nuc. **F.** As in D but NS1-nuc and HBov NS1-nuc.

### Prediction of nick site ssDNA recognition and cleavage by NS1-nuc

A structure of a more distantly related viral replication enzyme (PDB accession code 6WE1^*45*^, from Wheat Dwarf Virus, WDV)^*46*^ provides the basis for a model of sequence specific ssDNA recognition and nicking by NS1-nuc. The C_α_ atoms of the active site residues (His81, His83, Tyr141 in Form I NS1-nuc and His59, His61, Phe106 in WDV Rep-nuc, which contains the active site mutation Y106F) were used to superimpose the two structures, and the position of the ssDNA bound to WDV Rep-nuc identified (yellow, **Fig. 5A**). A feature of the WDV Rep-nuc structure, the “ssDNA bridging motif”^*46*^ (orange, **Fig. 5A**), is directly involved in ssDNA binding, making numerous interactions to the bases of several nucleotides and bridging the 5’ and 3’ ends of the bound ssDNA. In NS1-nuc, an insert is found in this position (magenta, **Fig. 5A**), which is predicted to interact with dsDNA (discussed further below), but also likely interacts with ssDNA. **Figures S7A-B** show the electrostatic potential maps of NS1-nuc (mapped onto the protein surface in **Fig. S7A**, and as a field in **Fig. S7B**), where blue indicates high positive charge, and red indicates high negative charge. The predicted ssDNA binding face (shown by the position of ssDNA taken from the alignment with the WDV Rep-nuc structure, magenta in **Fig. S7A**, yellow in **Fig. S7B**), shows a high degree of positive charge consistent with binding to negatively charged DNA. Also consistent with this predicted DNA binding face are the positions of the bound phosphate ions (orange and red spheres, **Fig. 5B**, magenta sticks, **Fig. 6**).

**Figure 5.**
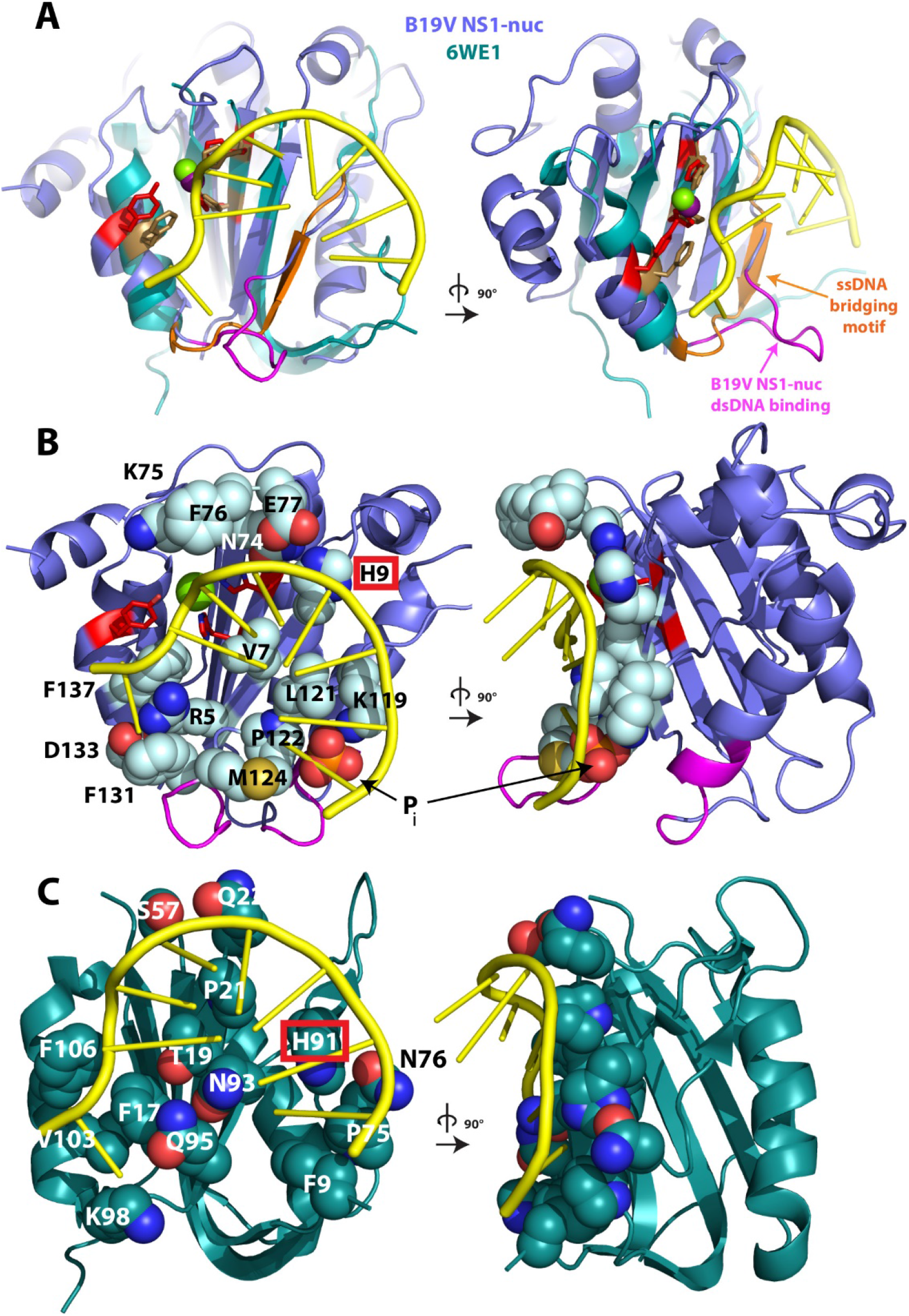
Nicking site recognition in ssDNA by WDV Rep-nuc and NS1-nuc. **A.** Orthogonal views of Form I NS1-nuc (slate blue) overlayed with a structure of Wheat Dwarf Virus (WDV) Rep-nuc (teal, PDB accession code 6WE1^*45*^) bound to ssDNA (yellow). Mg^2+^ from Form II NS1-nuc shown as green sphere, Mn^2+^ from WDV Rep-nuc shown as dark purple sphere. Active site residues of NS1-nuc and WDV Rep-nuc are shown in red and brown, respectively. The “ssDNA bridging” segment of WDV Rep-nuc is shown in orange, and residues 123-132 of NS1-nuc (a putative dsDNA binding element) shown in magenta. **B.** As in A, but with residues implicated in ssDNA binding shown as spheres. Red box indicates the residue H9, which appears to be in a similar position as a H91 in WDV Rep-nuc. **C.** Orthogonal views of the nuclease domain of WDV Rep-nuc bound to nick site ssDNA (PDB accession code 6WE1^*45*^) with residues within hydrogen bonding, van der Waals, or salt bridging distance shown as spheres. Red box indicates the residue H91, which appears to be in a similar position as a H9 in B19V NS1-nuc.

**Figure 6.**
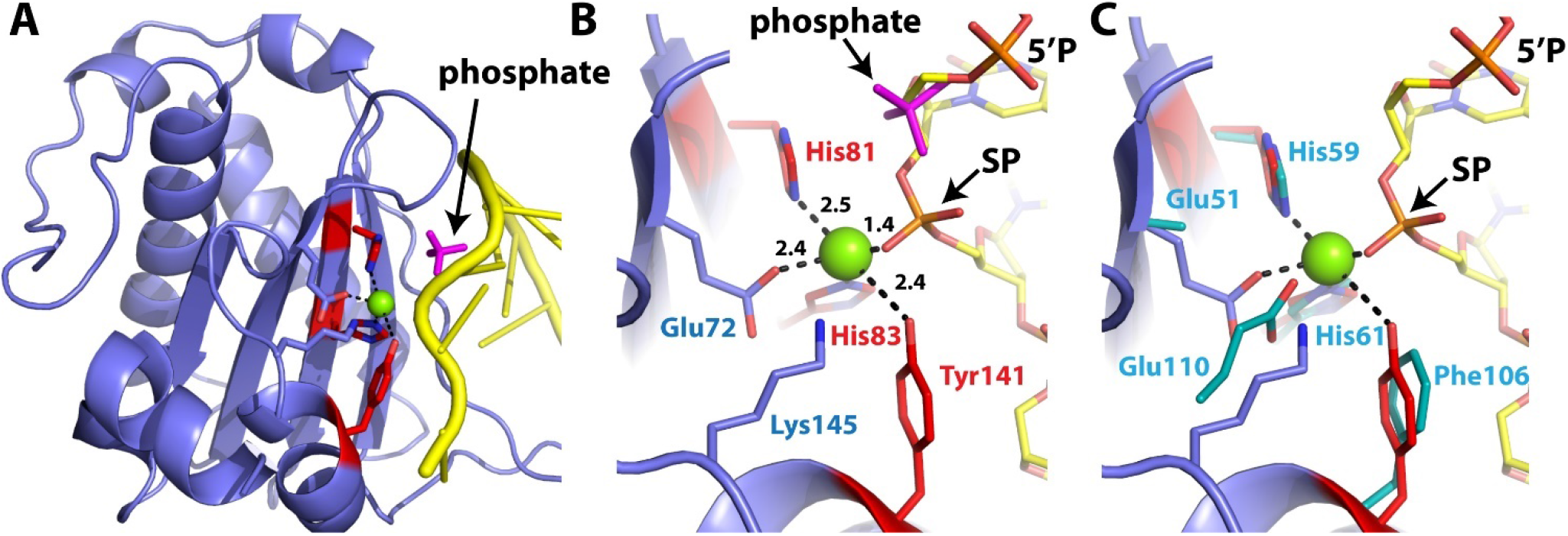
Phosphate bound near NS1-nuc active site. **A.** Form I NS1-nuc shown in cartoon, with active site residues in stick form (red and blue). Mg^2+^ from Form II shown as a green sphere. ssDNA (shown in yellow) from WDV Rep-nuc/ssDNA (PDB accession code 6WE1^*45*^) after superposition on to Form I NS1-nuc using the C_α_ of the two His and Tyr active site residues. The bound phosphate of Form I is shown in magenta sticks, and closely approaches the backbone of the modeled ssDNA. **B.** As in A, with DNA shown in stick form, colored by atom type. The phosphorus atom of the bound phosphate ion is 3.5 Å from the scissile phosphate (SP) phosphorus atom and 3.6 Å from the 5’ neighboring phosphate (5’P) phosphorus atom. Distances between Mg^2+^ and atoms of His81, His83, Tyr141, Lys145, Glu72, and a non-esterified oxygen of the scissile phosphodiester shown in Å. **C.** As in B, with selected side chains of WDV Rep-nuc shown in teal.

The model of NS1-nuc bound to ssDNA created from the superposition with the WDV Rep-nuc structure allows for the prediction of residues likely to be involved in ssDNA recognition (light blue, red, and dark blue spheres, **Fig. 5B**). Only His9 of NS1-nuc (H9, red boxes, **Fig. 5B**) and His91 of WDV Rep-nuc (H91, **Fig. 5C**) appear to be conserved between the two structures. The model of NS1-nuc bound to nick site ssDNA also provides for the opportunity to model atoms in the active site **(Figs. 6-7**). Both WDV Rep-nuc and NS1-nuc are members of the HUH nuclease superfamily, which require a divalent cation for DNA cleavage activity^*47*^. Prior work with NS1-nuc showed that Mg^2+^, Co^2+^, Ni^2+^, and Mn^2+^ but not Zn^2+^, Ca^2+^, or Cu^2+^ confer DNA cleavage activity. Both the Mg^2+^ (from Form II NS1-nuc) and Zn^2+^ (from NS1-nuc in PDB accession code 6USM^*31*^) are bound in the same position in the active site, therefore the difference in activity with these two ions may derive from different chemical properties such as ligation geometries and ability to polarize ligated atoms^*48–50*^. In the current model of NS1-nuc bound to ssDNA, the Mg^2+^ (green sphere, **Fig. 6B**) is positioned near a non-esterified oxygen of the scissile phosphate (SP, **Fig. 6B-C**, the bond to be cleaved in the nicking/nuclease reaction). The distance between the modeled phosphate oxygen and Mg^2+^ is 1.4 Å, somewhat closer than the typical ligation distance for Mg^2+^ to oxygen ligands (1.9-2.1 Å)^*51*^, but small adjustments in the position of the bound DNA could easily bring this distance to a more optimal value. The Mg^2+^ is also within ligation distance of the side chains of the active site residues His81 (2.5 Å) and Tyr141(2.4 Å)(**Fig. 6B**), as well as that of Glu72 (2.4 Å), and near His83 (3.3Å)(the relatively longer ligation distances may be due to coordinate error in the relatively low resolution of the Form II structure). WDV Rep-nuc also possesses a nearby glutamic acid residue in the active site capable of ligation to the Mg^2+^ (Glu110, **Fig. 6C**). Divalent cation-dependent nucleases catalyze DNA cleavage by any or all of the following: 1) polarization of the nucleophile (in this case, the phenolic oxygen of Tyr141) to increase its nucleophilicity (this often occurs via ligation to the divalent cation and may result in deprotonation of the nucleophile), 2) stabilization of the transition state after nucleophilic attack (often via divalent cation ligation to a non-esterified oxygen of the scissile phosphate), and 3) stabilization of the leaving group (the O3’) following bond breakage (often via direct ligation to the divalent cation or protonation from a divalent cation ligated water molecule)^*45 52–54*^. In our NS1-nuc structure with modeled ssDNA, we find that the Tyr141 OH is within ligation distance to the Mg^2+^ (2.4 Å), and the Mg^2+^ is within ligation distance to one non-esterified oxygen of the scissile phosphate, hence fulfilling two of the predicted catalytic configurations. In the case of stabilization of the leaving group, the ssDNA bound model does not predict a direct ligation of the O3’ leaving group to the Mg^2+^, but protonation by a Mg^2+^ ligated water molecule could be possible. Divalent cation dependent nucleases also accelerate DNA cleavage by organizing reactive groups in the active site into a geometry favorable for nucleophilic attack and bond breakage^*45, 52–54*^. This geometry includes: 1) positioning the nucleophile within van der Waals radii of the phosphorus atom (the atom to be attacked by the nucleophile), and 2) arranging the three atoms of the bond making and breaking reaction (the attacking group, the phosphorus atom, and the leaving group) in an “in-line” configuration such that the angle between them is 180°^*54*^. We find in our model that the Tyr141 hydroxyl oxygen atom (the nucleophile of the DNA nicking reaction) is 3.3 Å from the phosphorus atom of the phosphodiester to be cleaved (the estimated van der Waals radii of the two atoms is 3.3 Å, 1.5 Å for oxygen, and 1.8 Å for phosphorus^*55*^), and the angle between the nucleophile, phosphorus atom, and leaving group (O3’ of the 5’ nucleotide) is 149°. Hence the active site moieties are poised in an appropriate position for the catalytic reaction to occur. Finally, the terminal amine of the Lys145 side chain is within hydrogen bonding distance to the Tyr141 OH nucleophile, suggesting a contribution to the catalytic reaction by acting as a general base to accept a proton from the nucleophile, or alternatively to stabilize a negative charge on the nucleophile following proton loss.

**Figure 7.**
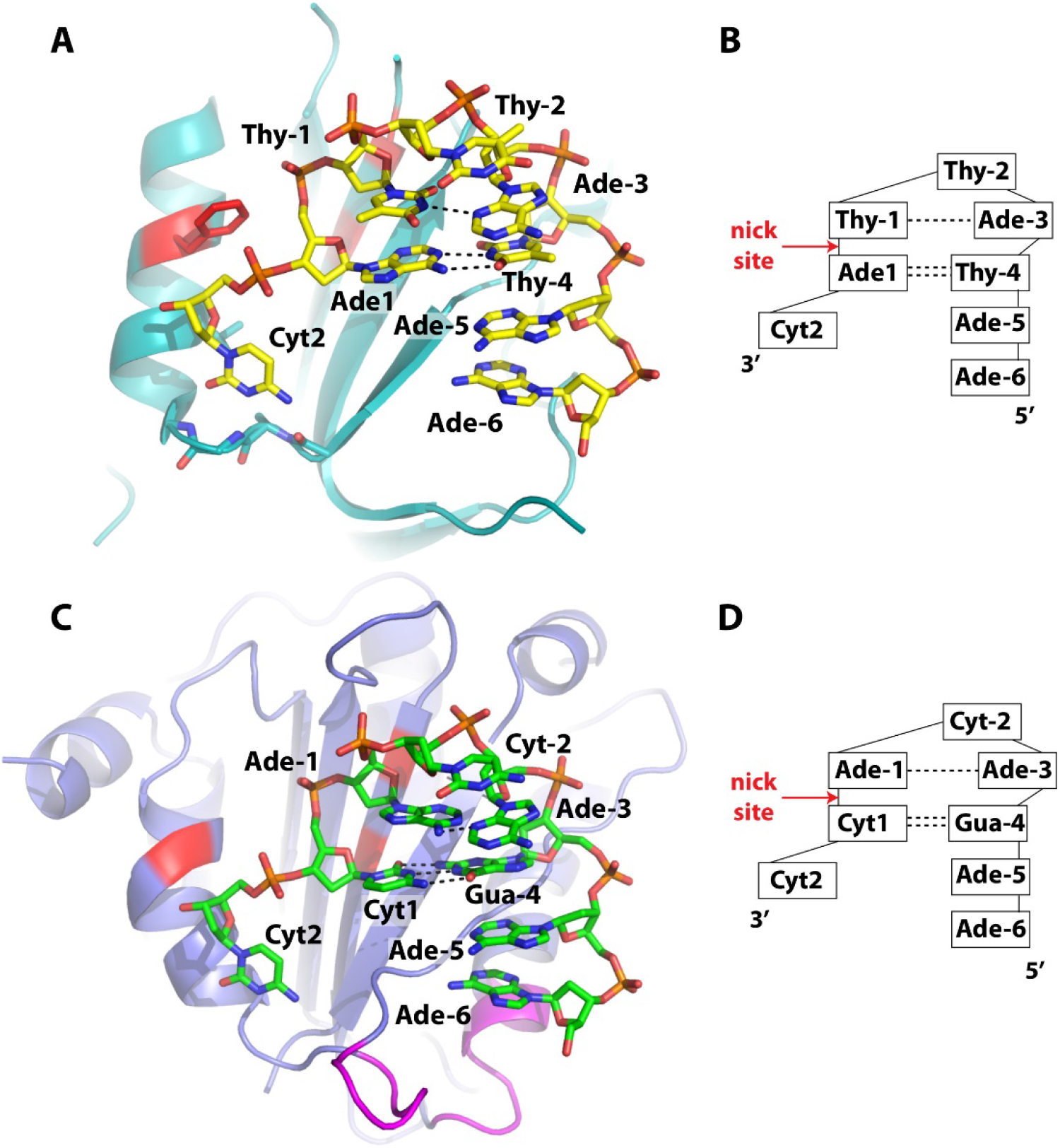
Nick site DNA binding by WDV Rep-nuc and NS1-nuc. **A.** Wheat Dwarf Virus (WDV) Rep-nuc bound to ssDNA containing the nick site (PDB accession code 6WE1^*45*^). **B.** Sequence of DNA in the WDV origin of replication around the nick site. Dashed lines indicate hydrogen bonding between bases. Stacking of bases is indicated by vertical alignment of bases. **C.** Model of NS1-nuc bound to nick (trs) site DNA based on the WDV Rep-nuc/DNA structure shown in A. **D.** B19V nick site DNA sequence and predicted interactions between bases.

The structure of WDV Rep-nuc bound to nick site DNA (**Fig. 7A**) suggests that sequence specific recognition occurs through a combination of direct readout, consisting of hydrogen bonds and van der Waals interactions to the chemically distinct portions of the DNA bases, as well as indirect readout, derived from the sequence specific energetics of DNA structure and base stacking^*46*^. Direct readout of the nick site DNA sequence is accomplished via hydrogen bonds between residues of WDV Rep-nuc and bases of nucleotides at the 2, −5, and −6 positions (see nucleotide numbering in **Fig. 7B**)^*46*^. Indirect readout of the nick site DNA sequence is suggested by the distorted U-shape of the bound ssDNA, as well as the base pairs between Ade1 and Thy-4 (a Watson-Crick bp) and Thy-1 and Ade-3 (a non-Watson-Crick bp). To understand how NS1-nuc recognizes its nick site in ssDNA, we substituted the DNA sequence of the ssDNA in the NS1-nuc/ssDNA model derived from the superposition with the WDV Rep-nuc/ssDNA structure (**Fig. 7C-D**). First, in terms of direct readout, the low amino acid sequence conservation in the DNA binding site residues makes direct readout contacts difficult to predict between NS1-nuc and the bound ssDNA, with the possible exception of the 2 position of the nick site DNA (which is Cyt in both viral nick sites, **Figs. 7B,D**). WDV Rep-nuc recognizes this base with hydrogen bonds from protein backbone atoms to the base pairing atoms Cyt2 (**Fig. 8A**). In the model of NS1-nuc bound to nick site DNA, hydrogen bonds between the side chain of Asp133 and the the O2 and N3 of Cyt2 would be possible with small adjustments (distances are 2.0 Å and 3.0 Å from the Asp133 carboxylate atoms to the O2 and N3 Cyt2 atoms, respectively)(**Fig. 8B**), predicting a role for Asp133 in sequence specific recognition at the 2 position of the nick site DNA. In addition, Arg5 approaches Cyt2 from behind the base, and Phe131 may form a π-hydrogen bond^*56*^ to the NH2 at the 4 of Cyt2 in this model, suggesting that these residues may also be important in DNA nick site sequence discrimination (**Fig. 8B**). In terms of indirect readout, the Watson-Crick bp between nucleotides at the 1 and −4 positions (Ade1 and Thy-4 in the WDV sequence, **Fig. 8C**) may be conserved as these nucleotides are Cyt1 and Gua-4 in the B19V nick site sequence (see model in **Fig. 8D**). The non-Watson-Crick base pair between Thy-1 and Ade-3 in the WDV structure occurs with a single hydrogen bond between the N3 of Thy-1 and N3 of Ade-3 (**Fig. 8E**). In the B19V nick site sequence, these bases are Ade-1 and Ade-3, and the structural model predicts that a single hydrogen bond is possible (after some adjustment due to the larger size of of the Ade base at the −1 position) between the N6 of Ade-1 and N3 of Ade-3 (green and dark blue sticks, **Fig. 8F**). Finally, in both models, a pyrimidine base is found at the turn of the bound DNA (Thy-2 in WDV, **Fig. 7A-B**, and Cyt-2 in B19V, **Fig. 7C-D**), which may be an important factor in indirect readout of the DNA sequence due to the formation of stacking interactions with the nucleotide at position −3 (an Ade in both cases).

**Figure 8.**
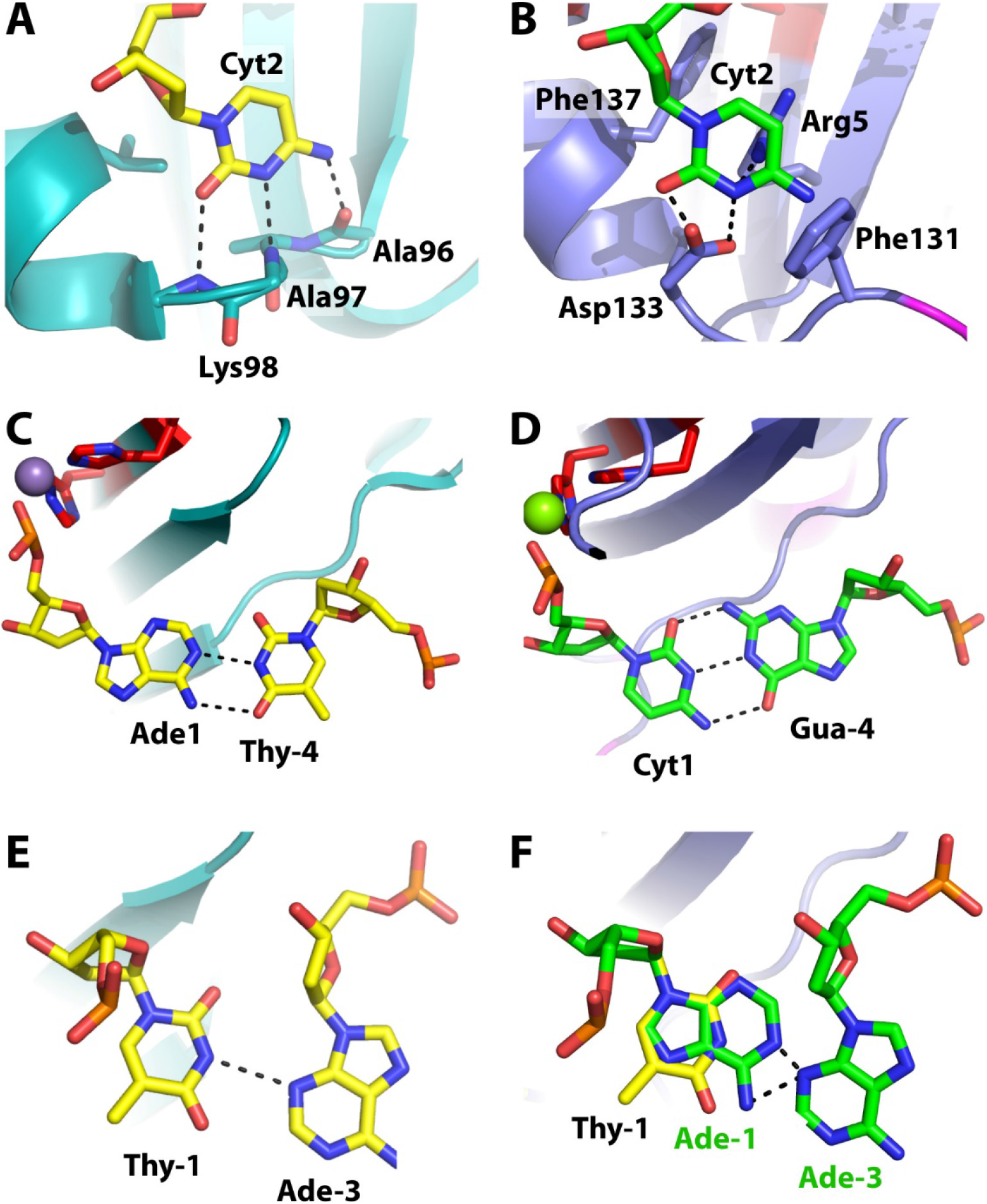
ssDNA sequence recognition by WDV Rep-nuc and predicted interactions between NS1-nuc and ssDNA. **A.** Interactions between Cyt2 and WDV Rep-nuc (PDB accession code 6WE1^*45*^). **B.** Model for interactions between Cyt2 and NS1-nuc. **C.** Watson-Crick base pair formed between Ade1 and Thy-4 formed in DNA bound by WDV Rep-nuc. Active site residues are shown in red and blue stick, and bound Mn^2+^ shown as purple sphere. **D.** Predicted Watson-Crick base pair formed by Cyt1 and Gua-4 in the model of B19V nick (trs) site DNA bound to NS1-nuc. **E**. Non-Watson-Crick base pair in WDV Rep-nuc bound DNA between Thy-1 and Ade-3. **F.** B19V NS1 nick (trs) site contains Ade at −1 and Ade at −3. Green shows interactions between −1 and −3 predicted in the model of NS1-nuc bound to nick site ssDNA, yellow shows the position of the Thy-1 in the structure with WDV Rep-nuc bound to DNA (coordinates for Ade-3 of the WDV structure overlap with the B19V model). Dashed lines indicate close approach of N6 of Ade-1 and N3 of Ade-3 (2.3 Å). Adjustments in positioning of bases would be necessary to bring this to a reasonable distance, such as 2.8 Å.

A study of sequence preferences at the B19V nick site was performed with the NS1-nuc domain^*23*^, which found the greatest preferences at nucleotides Cyt2, Cyt1, and Ade-1, with Cyt2 being most important, followed by Cyt1 and Ade-1. The nucleotides at these positions were also found to be the most important for nicking by WDV Rep-nuc using a sequence specificity selection approach (HUH-seq)^*46*^, and in the same order of importance. In the case of B19V, exhaustive substitutions of the nick site sequence were not tested, instead only mutation to the Watson-Crick base pairing partner was examined for cleavage by NS1-nuc. However, the large decrease in DNA cleavage activity of NS1-nuc found as a result of changing Cyt2 to a Gua^*23*^ may be explained by the predicted tight binding pocket surrounding Cyt2 (**Fig. 8B**), since a Gua base would be too large to fit into this pocket and could not form the same interactions with Asp133 and Phe131. The next most significant decrease in cleavage activity occurred with the substitution of Cyt1 to Gua. Since the model predicts a base pairing interaction with Gua-4 (**Fig. 8D**), it is clear that substitution of the Cyt with a Gua would disrupt this pairing. Strangely, substitution of Gua-4 with Cyt resulted in an increase in DNA cleavage by NS1-nuc. The Cyt-Cyt interaction (between Cyt1-Cyt-4), while not predicted to be favorable, may result in less disruption to the structure of the bound DNA due to the relatively small size of the Cyt bases (compared to two Gua bases as in Gua1-Gua-4). Finally, substitution of Ade-1 with Thy diminished cleavage by approximately 50%^*23*^. This substitution would disrupt the predicted hydrogen bonding with Ade-3, between the N6 of Ade-1 and the N3 of Ade-3 (green and blue sticks, **Fig. 8F**). A Thy at position −1 would not offer a hydrogen bond donor to take the place of the N6 of Ade-1, but interestingly, is the sequence seen in the WDV structure (Thy-1, yellow, **Fig. 8F).**

The fact that substitution of Ade-1 with Thy is detrimental to DNA cleavage by NS1-nuc suggests a different configuration of these bases in the two structures, which would be necessary to accommodate the wild type Ade-1-Ade-3 interaction predicted in the bound B19V ssDNA. Conversely, substitution of Ade-3 to Thy had little effect, possibly due to the availability of the O2 of a Thy to take the place of the N3 of Ade-3 in the predicted hydrogen bonding interaction (**Fig. 8F**). It may also be that substitutions near the nicking site (in −1, 1, and 2) are more sensitive due to the requirement for precise positioning of atoms in the active site in order to achieve optimal cleavage activity.

### Prediction of dsDNA binding by NS1-nuc

In addition to binding to ssDNA, B19V NS1 must also bind to target sequences in dsDNA known as NSBE, for NS1 Binding Elements. These sequences are located near the nick site (*i.e*. the trs) (**Fig. 1B**). It is the N-terminal nuclease domain of NS1 (*i.e*. NS1-nuc) which is also responsible for recognition of these sequences^*23*^. The structure of the nuclease domain from AAV Rep (AAV Rep-nuc) bound to dsDNA containing RBE repeats (PDB accession code 1RZ9^*57*^), which are sequences analogous to the NSBE of B19V, provides a framework for modeling the interactions of NS1-nuc with the NSBE sequences in dsDNA. **Figure S8A** shows a map of the AAV Rep-nuc domains bound to the RBE sites in dsDNA in this crystal structure. Five copies of AAV Rep-nuc (colored ovals, **Fig. S8A**) bind the five four-base (imperfect) RBE repeats (boxed bp, **Fig. S8A**), and follow the helical twist of the DNA since each copy of AAV Rep-nuc interacts with bases in both the major and minor grooves in an equivalent manner (**Fig. S8B**). Interactions to the DNA by each copy of AAV Rep-nuc are not confined to a single quartet, but instead overlap. For example, one copy (yellow, **Fig. S8A**) interacts with base pairs of the first RBE quartet (boxed sequences, counting from left to right, **Fig. S8A**) in the major groove, as well as base pairs in the minor groove in the second RBE quartet. The next AAV Rep-nuc copy (green, **Fig. S8A**) interacts with the second RBE quartet bases via the major groove, and third quartet bases via the minor groove. Each AAV Rep-nuc copy therefore interacts with 2 quartets, and each quartet interacts with two copies of AAV Rep-nuc. **Figure S8C** shows the NSBE and trs sequences of the B19V origin of replication. Rather than five quartet repeats, the sequence appears to have four octet repeats. The octet sequences are very GC rich, and the first two boxed octets are identical in DNA sequence. Creating a model for NS1-nuc binding to DNA with this repeat pattern, based on the structure of AAV Rep-nuc bound to RBE containing DNA, is difficult given the different repeat sequences, lengths, and spacings, as well as the high amino acid sequence divergence between AAV Rep-nuc and NS1-nuc. However, it is possible that NS1-nuc binds the NSBE sequences in a manner similar to that of AAV Rep-nuc, using overlapping quartets rather than four separate octets. Prior investigations indicate that 5-7 copies of NS1-nuc bind to the DNA containing all four NSBE sequences, with those of the first three octets being the most important for binding. These sequences amount to 28 bp, or 7 quartets, consistent with 7 copies of NS1-nuc binding in the AAV Rep-nuc pattern. However, more information on NS1 interactions with dsDNA containing the NSBE sequences will be necessary to assign the exact positioning and amino acid-nucleotide interactions between NS1-nuc and the DNA sequence.

Although precise protein-DNA contacts between NS1-nuc and the NSBE sequences are not possible to predict as of yet, a model based on the AAV Rep-nuc structure does allow for prediction of likely dsDNA binding residues in the amino acid sequence of NS1-nuc. Two segments of AAV Rep-nuc interact with dsDNA, and these same segments are largely conserved in the structure (but not sequence) of NS1-nuc. **Figures 9A-B** show structural alignments of NS1-nuc (blue) on AAV Rep-nuc (yellow), with suspected dsDNA interacting residues of NS1-nuc in magenta. **Figure 9C** shows an alignment of residues in the two dsDNA binding loops, with the dsDNA bound to AAV shown in cartoon form, identifying the locations of the loop which interacts predominantly with the dsDNA minor groove (101-111 in AAV Rep-nuc, 93-103 in NS1-nuc), and residues interacting with the major groove (135-142 in AAV Rep-nuc, 124-130 in NS1-nuc). The phosphate ion bound to NS1-nuc the Form I structure is located very near the predicted binding site for dsDNA (orange and red spheres, **Fig. 9D**). Thus, we find that the phosphate ions bound to NS1-nuc are located in both the ssDNA (in two separate positions) and dsDNA binding clefts, suggesting a strong affinity for phosphate moieties such as those in DNA in these locations and lending further support to these model for single- and double-stranded DNA binding.

**Figure 9.**
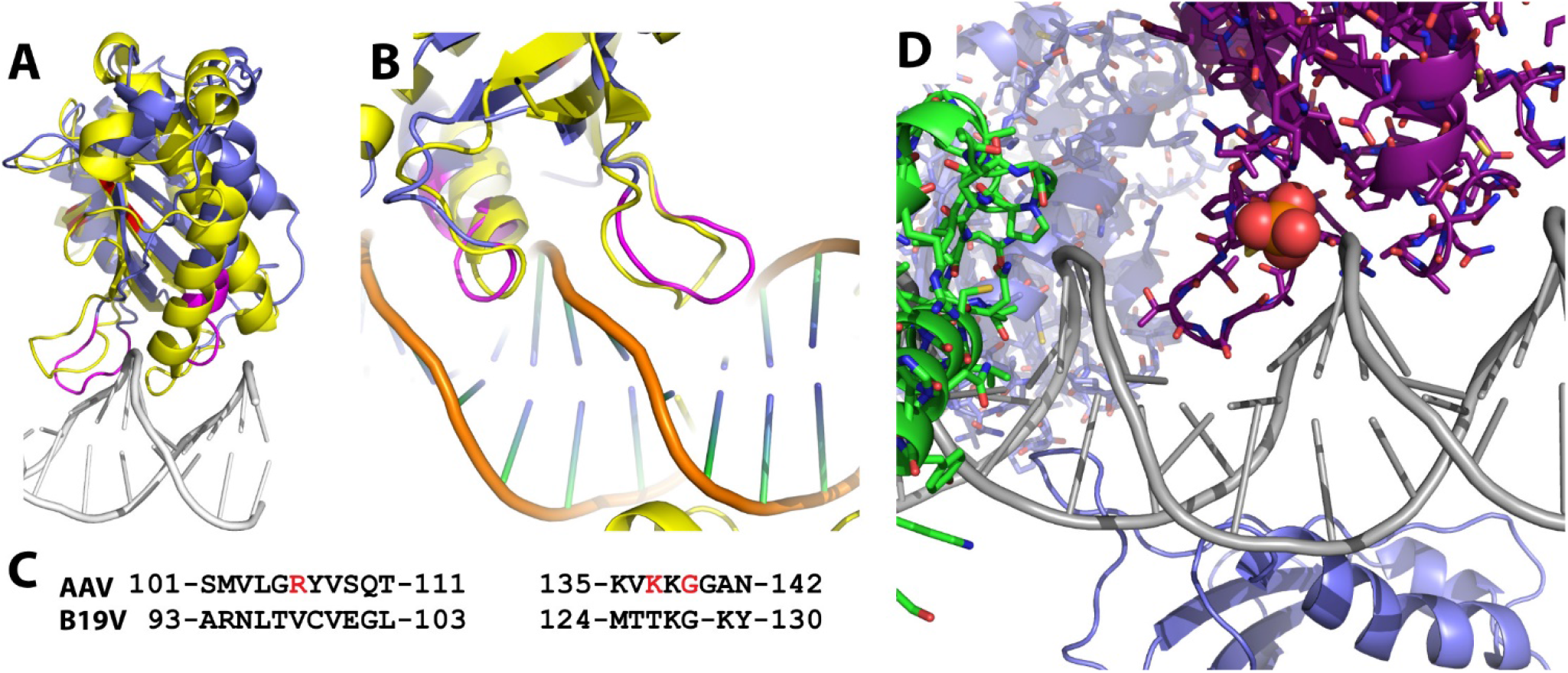
Predicted recognition of NSBE containing dsDNA by NS1-nuc. **A.** Overlay of NS1-nuc (blue) on AAV Rep-nuc (yellow) bound to RBE containing dsDNA (white). Putative DNA binding residues of NS1-nuc shown in magenta and active site residues highlighted in red. **B.** Close-up of B, with DNA cartoon shown in orange, green, and blue. **C.** Residues of the segments of AAV Rep-nuc closely approaching or interacting with the dsDNA (top sequence), and corresponding residues of NS1-nuc after least squares alignment of the two protein structures. Residues shown in red indicate sequence specific contacts from AAV Rep-nuc to the DNA (PDB accession code 1RZ9^*57*^) (top). **D.** The location of a bound phosphate ion (orange and red, spheres) in the Form I NS1-nuc structure is found near the predicted dsDNA binding site of the model shown in B.

Prior binding studies indicate that NS1-nuc binds DNA cooperatively, as evidenced by the shape of binding isotherms^*23*^. Cooperativity indicates that the binding of DNA by one copy of NS1-nuc increases the affinity of subsequent copies of NS1-nuc to the same DNA. This effect can occur when favorable protein-protein interactions occur between copies of NS1-nuc on the DNA, and/or when distortions made to the DNA upon binding of one copy facilitate the binding of subsequent copies by negating the requirement for the DNA distortion (which costs energy) by those subsequent protein copies. Cooperative DNA binding by AAV Rep-nuc is also suggested by the shape of the binding isotherm shown in Hickman, *et al*. (Figure 4B, lowest panel)^*58*^. However, no protein-protein interactions are found between copies of AAV Rep-nuc bound to dsDNA in the crystal structure, and similarly, no protein-protein interactions are predicted when the NS1-nuc structure is superimposed onto each of the five copies of AAV Rep-nuc bound to dsDNA (the closest approach between different copies of NS1-nuc is 8 Å, **Fig. S9**). It is possible that residues beyond the C-terminus of AAV Rep-nuc or NS1-nuc (such as in the linker which connects the nuclease domain to the central helicase domain) extend enough to allow contacts between neighboring copies of AAV Rep-nuc or NS1-nuc when bound to DNA. Indeed, when a structure of AAV Rep-nuc containing these linker residues (PDB accession code 5DCX)^*42, 59*^ is superimposed onto each of the five AAV Rep-nuc copies of the AAV Rep-nuc/dsDNA structure, the linker residues extend far enough to form interactions with neighboring AAV Rep-nuc copies (**Fig. S10**). However, these residues were not present in the prior binding studies which showed strongly cooperative DNA binding by both AAV Rep and B19V NS1 nuclease domains^*23, 58*^, suggesting the existence of some other mechanism of cooperativity beyond protein-protein interactions.

The other likely origin of cooperative DNA binding involves distortions in the bound DNA, where binding of each copy of a DNA binding protein distorts the DNA in a way which results in an increase in binding affinity of subsequent copies. Analysis of the DNA conformation in the AAV Rep-nuc/dsDNA structure indeed shows distortions from ideal B-form DNA, as was also discussed by the authors of this work^*58*^, and which show a pattern of distortions that follow the pattern of the quartet repeat (**Fig. S11**). Since binding sites of NS1-nuc to NSBE likely overlap as they do in RBE binding by AAV Rep-nuc, distortions induced by one copy of NS1-nuc could facilitate binding of neighboring copies by presenting DNA pre-distorted in the manner which complements the protein-DNA binding interface, and eliminates the cost of DNA distortion to the binding of that subsequent copy to the DNA. Hence, we predict based on this analysis that DNA distortions play a large role in the cooperativity seen in NSBE binding by NS1-nuc.

## Conclusions

Presented herein are two new structures of the N-terminal origin binding and DNA nicking domain of Human Parvovirus B19 NS1 protein (*i.e*. NS1-nuc). Phosphate ions are found bound to the Form I structure at predicted DNA binding sites including two sites in the predicted nick site ssDNA binding cleft, and one at the predicted dsDNA binding site. A second crystal form (Form II) solved at a lower resolution (3.5 Å) shows the position of a bound Mg^2+^ in the nicking active site, in nearly the same position as a Zn^2+^ found in a previously solved structure of NS1-nuc (PDB accession code 6USM^*31*^). A model for binding to ssDNA containing the nick site found at the origin of replication has been created and analyzed showing residues likely important in recognition of the nick site DNA sequence, as well as the arrangement of active site moieties in the DNA nicking active site. Positioning of the nick site ssDNA onto NS1-nuc was greatly facilitated by structural similarity to a distant homolog, WDV Rep-nuc (PDB accession code 6WE1^*45*^). Modeling based on the structural similarity between NS1-nuc and AAV Rep-nuc (PDB accession code 1RZ9^*57*^) allowed for identification of residues likely involved in major and minor groove contacts in dsDNA, however, exact positioning of NS1-nuc on the DNA sequences of the NSBE repeats was made difficult by the dissimilar nature of B19V NS1-nuc and AAV Rep-nuc sequences and the sequences in DNA they target. Further, the origin of cooperativity in binding of NS1-nuc to NSBE sequences in dsDNA is predicted to more likely originate from DNA distortions than from protein-protein contacts between adjacent copies of NS1-nuc bound to dsDNA. Experimentally determined structures of NS1-nuc bound to the single-stranded nick site and to the double-stranded NSBE sequences will be necessary to confirm or correct these existing models, and for precise positioning of NS1-nuc on the dsDNA.

## Abbreviations

AAV: Adeno Associated Virus
B19V: Human Parvovirus B19
NS1-nuc: B19V NS1 nuclease and origin binding domain
C_α_: alpha carbon atom
HBov: Human Bocavirus
MVM: Minute Virus of Mice
MVM NS1: NS1 protein from MVM
MVM NS1-nuc: the N-terminal nuclease and origin binding domain of MVM NS1
NS1: B19V NS1 unless otherwise indicated
NS1-nuc: the N-terminal nuclease and origin binding domain of B19V NS1
NSBE: B19V NS1 Binding Elements in double-stranded DNA at the B19V origin of replication
PAGE: polyacrylamide gel electrophoresis
AAV Rep: a homolog of NS1 from AAV
AAV Rep-nuc: the N-terminal endonuclease domain of AAV Rep
SDS: sodium dodecyl sulfate
TEV: Tobacco Etch Virus
Tris: Tris(hydroxymethyl)aminomethane
Tris-HCl: Tris titrated to a desired pH with HCl
trs: terminal resolution site or nicking site in DNA at the viral origin of replication
WDV: Wheat Dwarf Virus
WDV Rep-nuc: the N-terminal origin binding and nuclease domain of WDV Rep protein

## Acknowledgments

Research reported in this publication was supported by the National Science Foundation under Grant No. MCB-1934291 (to N.H.), the National Institute of General Medical Sciences of the National Institutes of Health via Grant T32GM008659 (to J.L.S.), and the University of Arizona Technology Research Initiative Fund (TRIF).Coordinates and structure factor amplitudes for Form I and II NS1-nuc structures have been deposited into the PDB under accession codes 7SZY and 7SZX.

## Supporting Information

### Sequences of constructs

#### A. Form I B19V NS1 nuclease domain (NS1-nuc) sequence

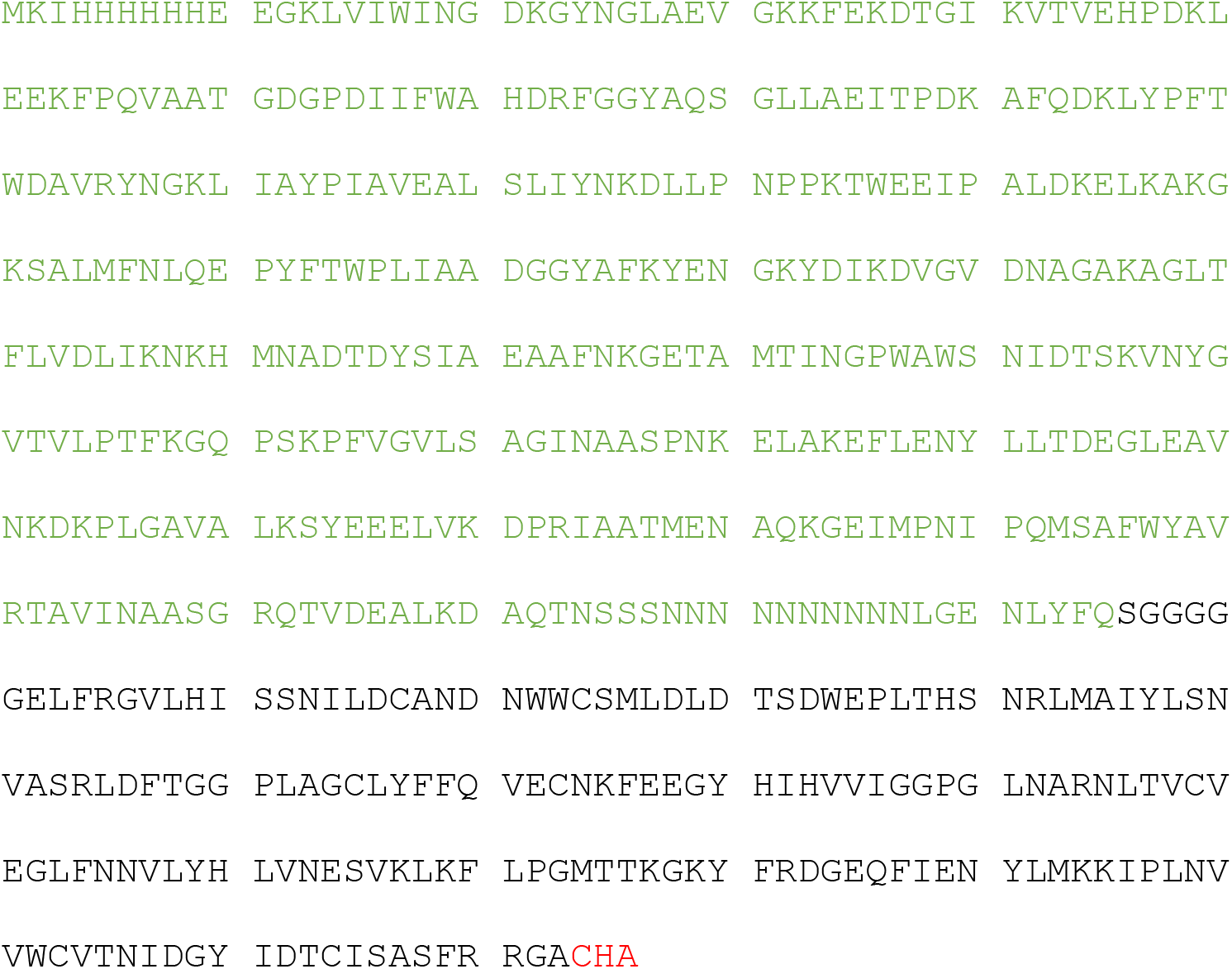

(Residues not ordered in the electron density map and not present in the PDB file are marked in red font, residues in green indicate purification tag residues not present in the final protein used in crystallization.)

### Numbering of the residues in Form I in the PDB file

Numbering matches that of the full length, wild type B19V NS1 protein, although the normal N-terminal M is replaced with a G residue in the recombinant protein:

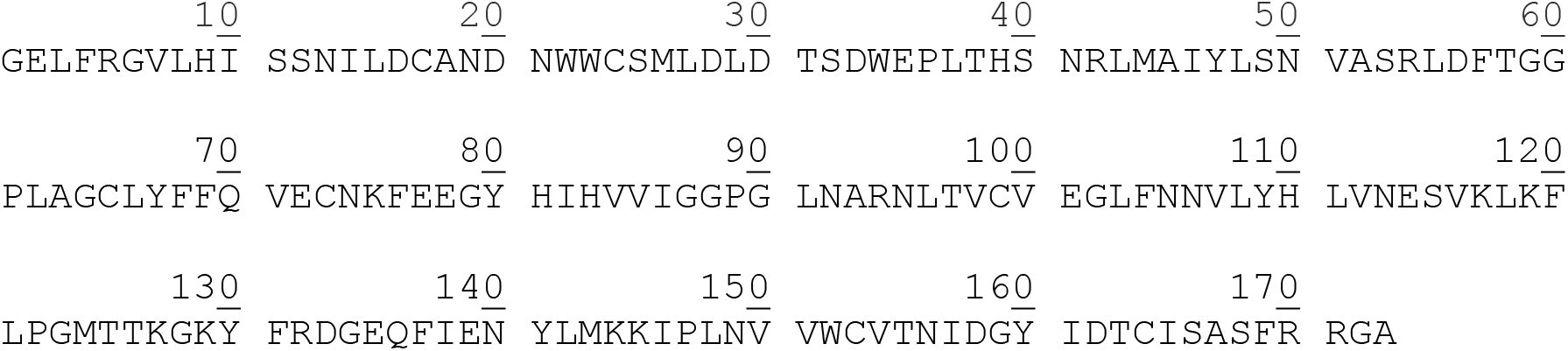

#### B. Form II B19V NS1-nuc

MSGSHHHHHHSSGENLYFQGELFRGVLHISSNILDCANDNWWCSMLDLDTSDWEPLTHSNRLMAIYLSNVASRLDFTGGPLAGCLYFFQVECNKFEEGYHIHVVIGGPGLNARNLTVCVEGLFNNVLYHLVNESVKLKFLPGMTTKGKYFRDGEQFIENYLMKKIPLNVVWCVTNIDGYIDTCISASFRRGACHAKKPRISANTDTVNNEGGESSCGGGDVVPFAGKG

(Residues not ordered in the electron density map and not present in the PDB file are marked in red font. Note that the full N-terminal tag was still present in the construct used in crystallization experiments.)

### Numbering of the residues in the PDB file (-indicates missing residue)

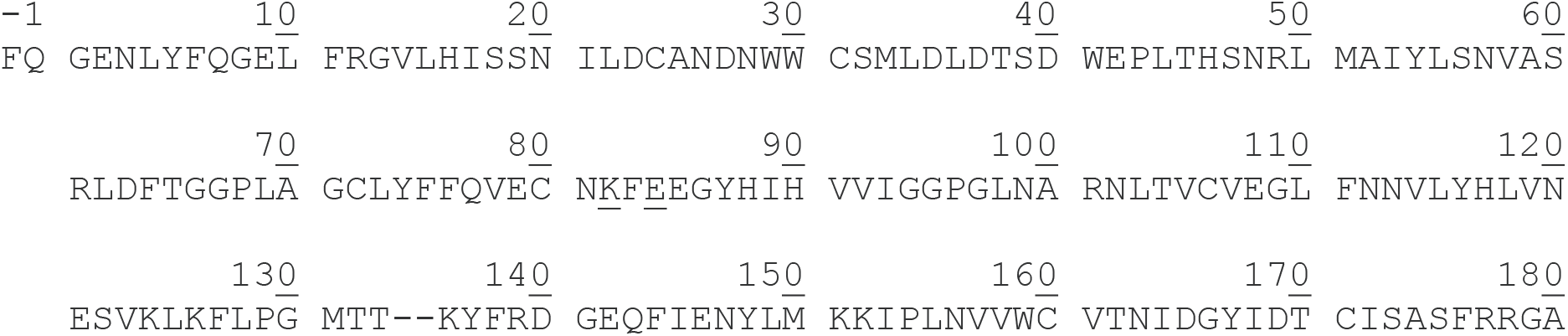

### DNA sequence used in crystallization drops of Form II NS1-nuc

NSBE1 and NSBE2 sequences highlighted in yellow:

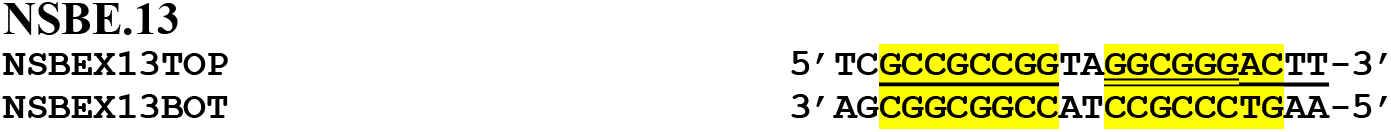

## Supplementary Figures

**Figure S1.**
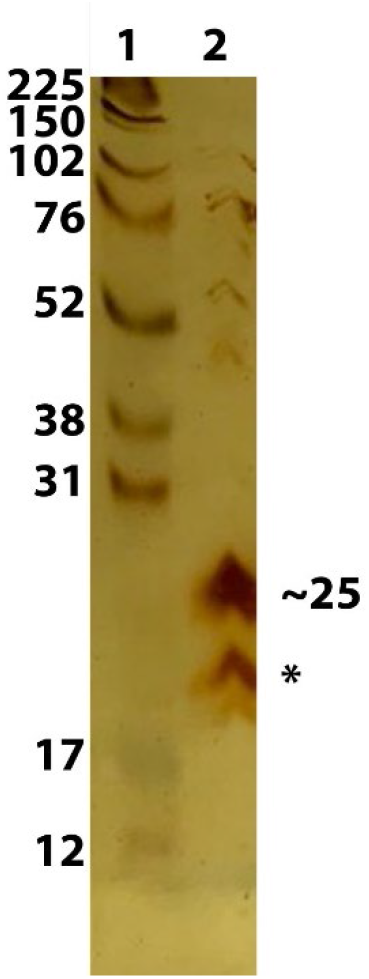
Composition of drops containing crystals of Form II NS1-nuc. Silver stained SDS-PAGE analysis of a drop containing Form II crystals (lane 2). The full length construct is predicted to be 25 kDa, however a smaller species indicated by the asterisk (*) may be the form found in the crystals.

**Figure S2.**
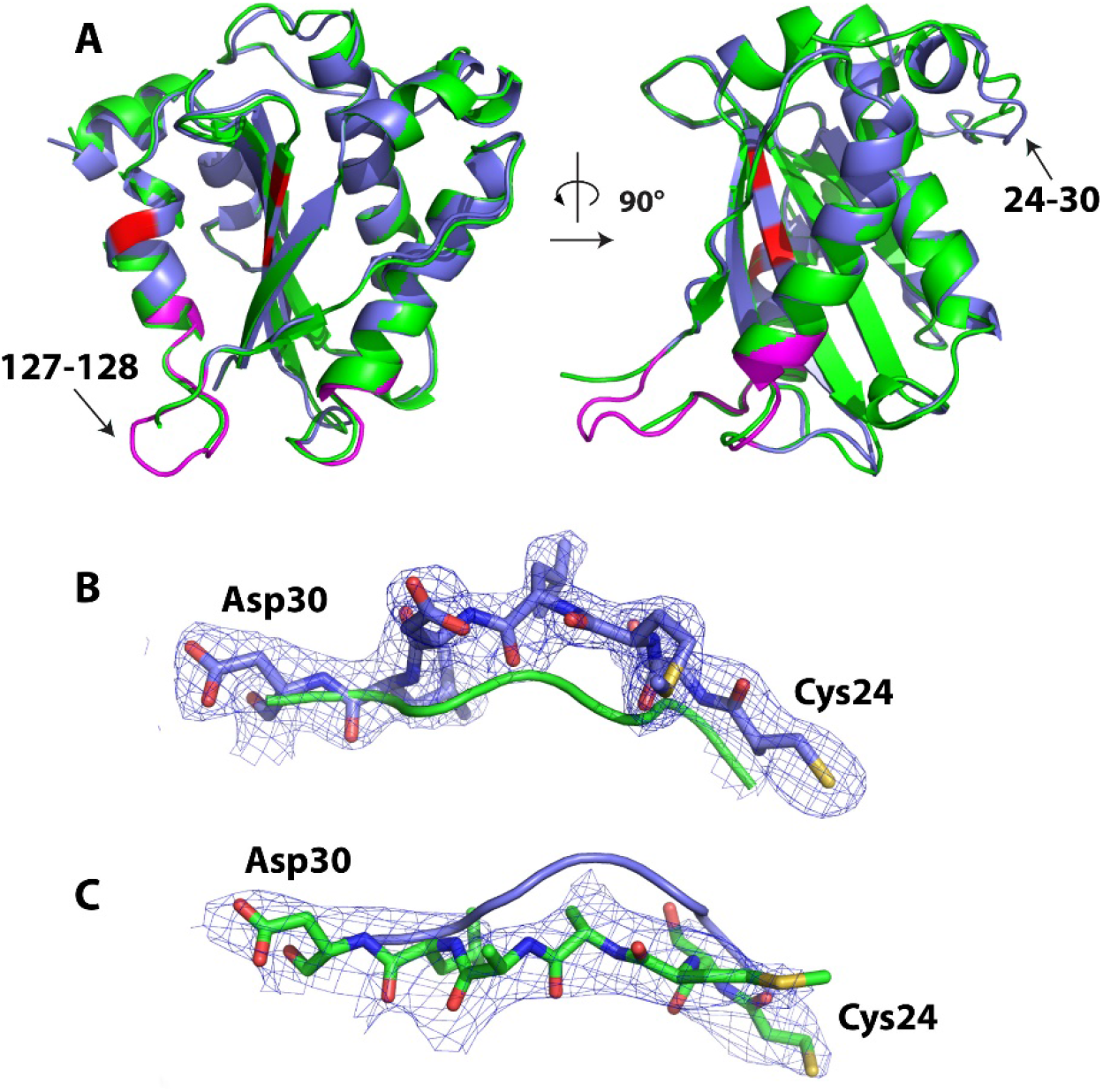
Comparison of NS1-nuc crystal structures from Forms I and II. **A.** Two views of a cartoon representation of Form I (blue) and Form II (green). Putative active site residues are shown in red, predicted dsDNA binding sites in Form I shown in magenta. Arrows indicate regions disordered in Form II (left), and a region (residues 24-30) of deviation in the mainchain position between the two forms (right). **B.** Form I 2Fo-Fc map (at 1σ) with residues 24-30 of the Form I model (blue, red, yellow). The mainchain trace of Form II is shown in green. **C.** Form II 2Fo-Fc map (at 1σ) with residue 24-30 of the Form II model (green, blue, red, yellow). The mainchain trace of Form I shown in blue.

**Figure S3.**
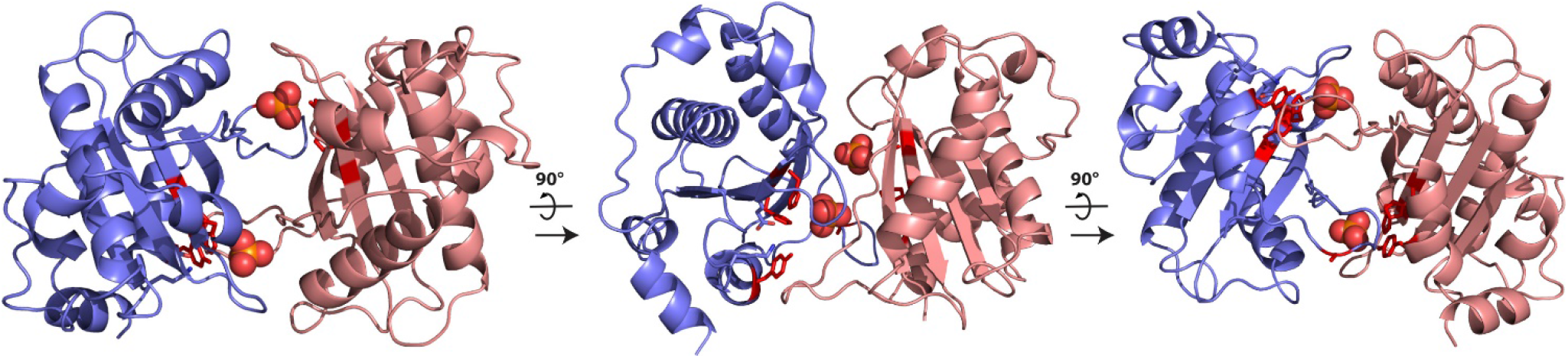
Two phosphate ions are bound between two copies of Form I NS1-nuc. One copy of NS1-nuc (slate blue) bound to one phosphate ion (orange and red spheres) form the asymmetric unit of Form I, however, a symmetry related copy (salmon) related by a crystallographic two-fold axis results in a second phosphate also positioned at the interface between the two copies of NS1-nuc. Active site residues shown as red sticks.

**Figure S4.**
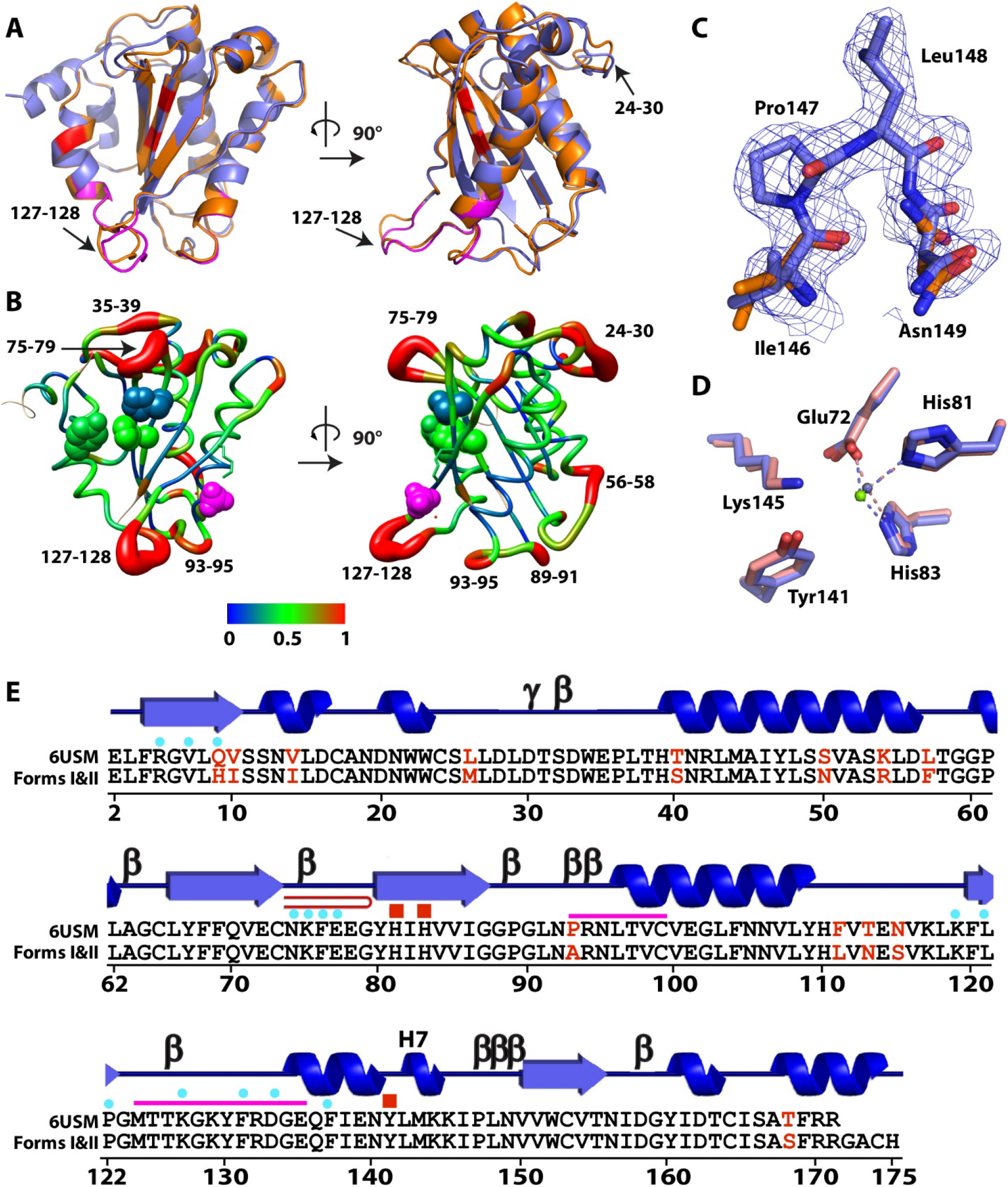
Differences in sequence and structure of NS1-nuc reported herein and previously (PDB accession code 6USM^*31*^). **A.** Orthogonal views of a superposition of the two structures (Form I structure in slate blue, that from 6USM in orange). Positions of active site residues are shown in red, and predicted dsDNA binding residues in magenta (Form I only). Regions of greater deviation identified with arrows. **B.** Orthogonal views of Form I NS1-nuc showing RMSD of C_α_ atoms between NS1-nuc and 6USM by color (see legend in Å). Residue numbers in segments with highest RMSD are also identified. The bound phosphate ion is shown as magenta spheres, and the active site residues (His81, His83, Tyr141) shown in spheres colored by C_α_ RMSD. **C.** 2Fo-Fc (1σ) map of Form I (in blue and red sticks) around the cis-proline at residue number 147. This region is not present in the model from 6USM (residues Ile146 and Asn149 shown in orange, blue, red sticks). **D.** Alignment of DNA nicking active sites from Form II (blue) and 6USM (light red). The modeled Mg^2+^ in Form II is shown in green, and the assigned Zn^2+^ of 6USM is shown in grey. **E.** Alignment of amino acid sequence of 6USM and Forms I and II crystal structures of NS1-nuc. Differences are highlighted in red text. Secondary structural elements are shown above the sequences, γ indicates γ-turn, β indicates β-turn, and red hairpin indicates β-hairpin. Active site residues are shown with red boxes above sequence, and predicted single- and double-stranded DNA binding residues are shown by blue dot or magenta line, respectively, above sequences.

**Figure S5.**
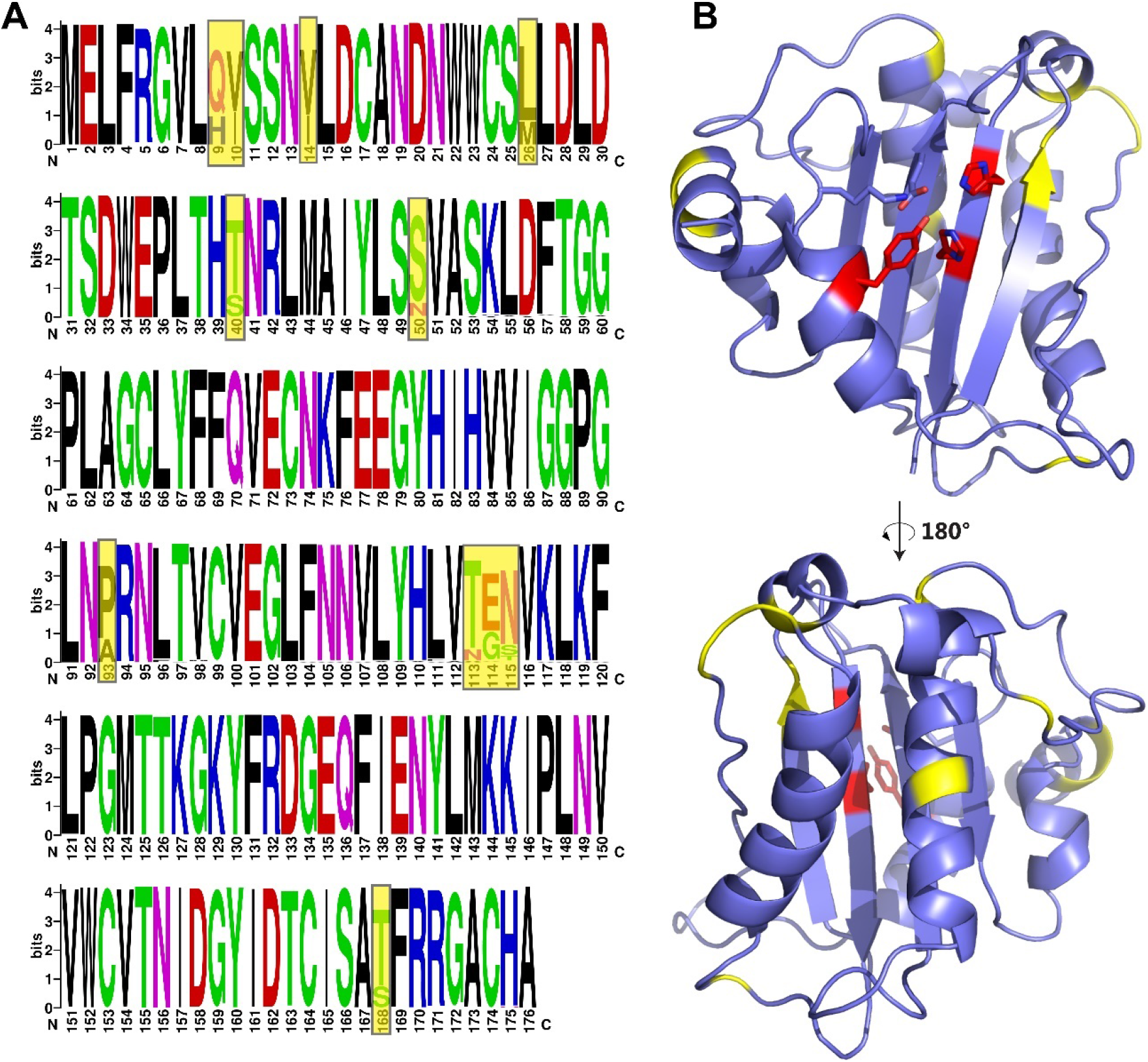
Sequence variations of NS1-nuc. **A.** 195 sequences of NS1-nuc (residues 1-176 of B19V NS1) were extracted from NCBI and the sequence variations shown in LOGO form^*41*^. Significant variations boxed in yellow. **B.** Two views of Form I NS1-nuc with positions of significant variants shown in yellow, and active site residues shown in red stick.

**Figure S6.**
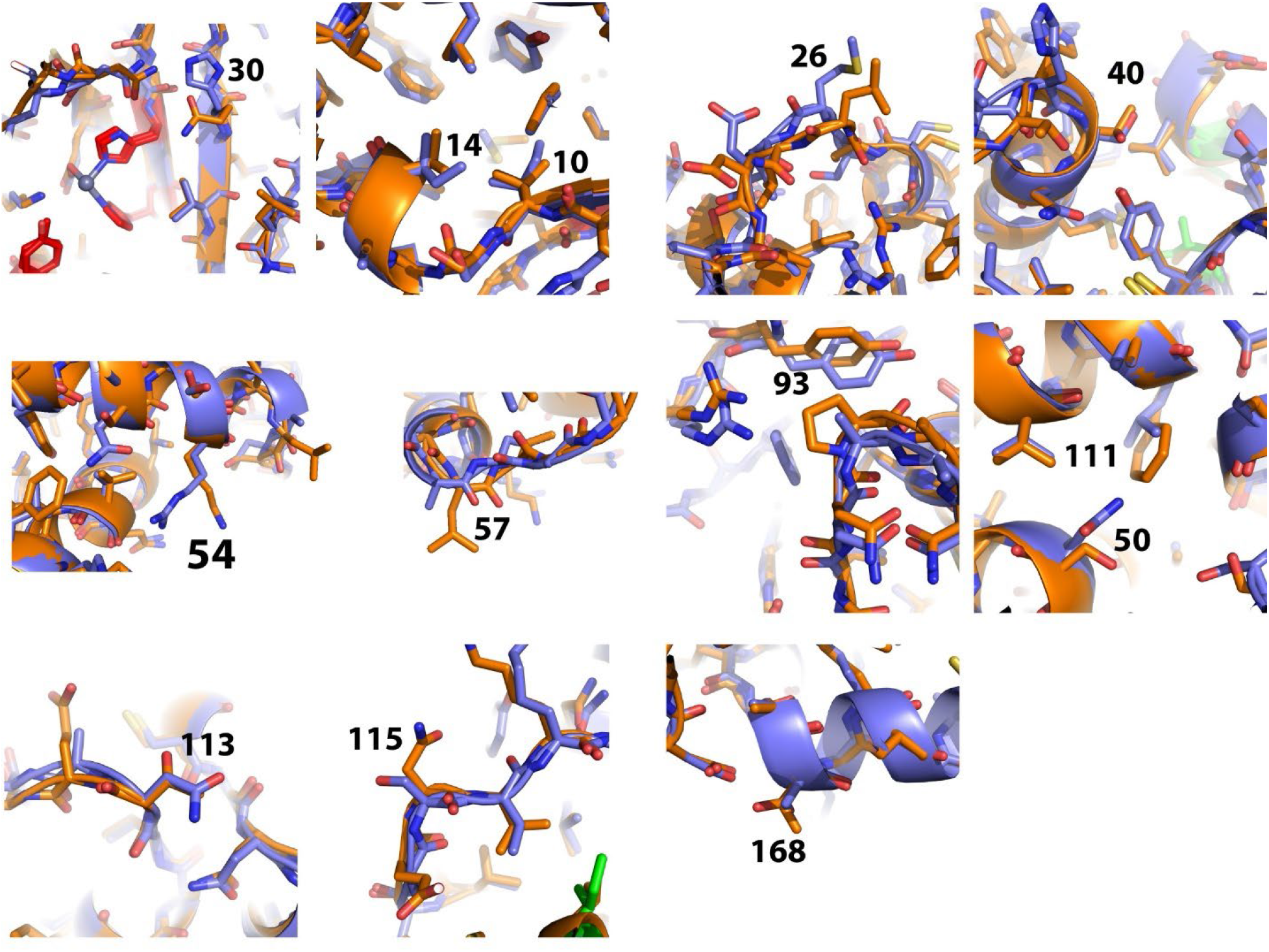
Structure around sequence variations in Form I and previously reported NS1-nuc. NS1-nuc (Form I) shown in slate blue, and NS1-nuc (PDB accession code 6USM^*31*^) shown in orange. Residue numbers of variants given in each panel

**Figure S7.**
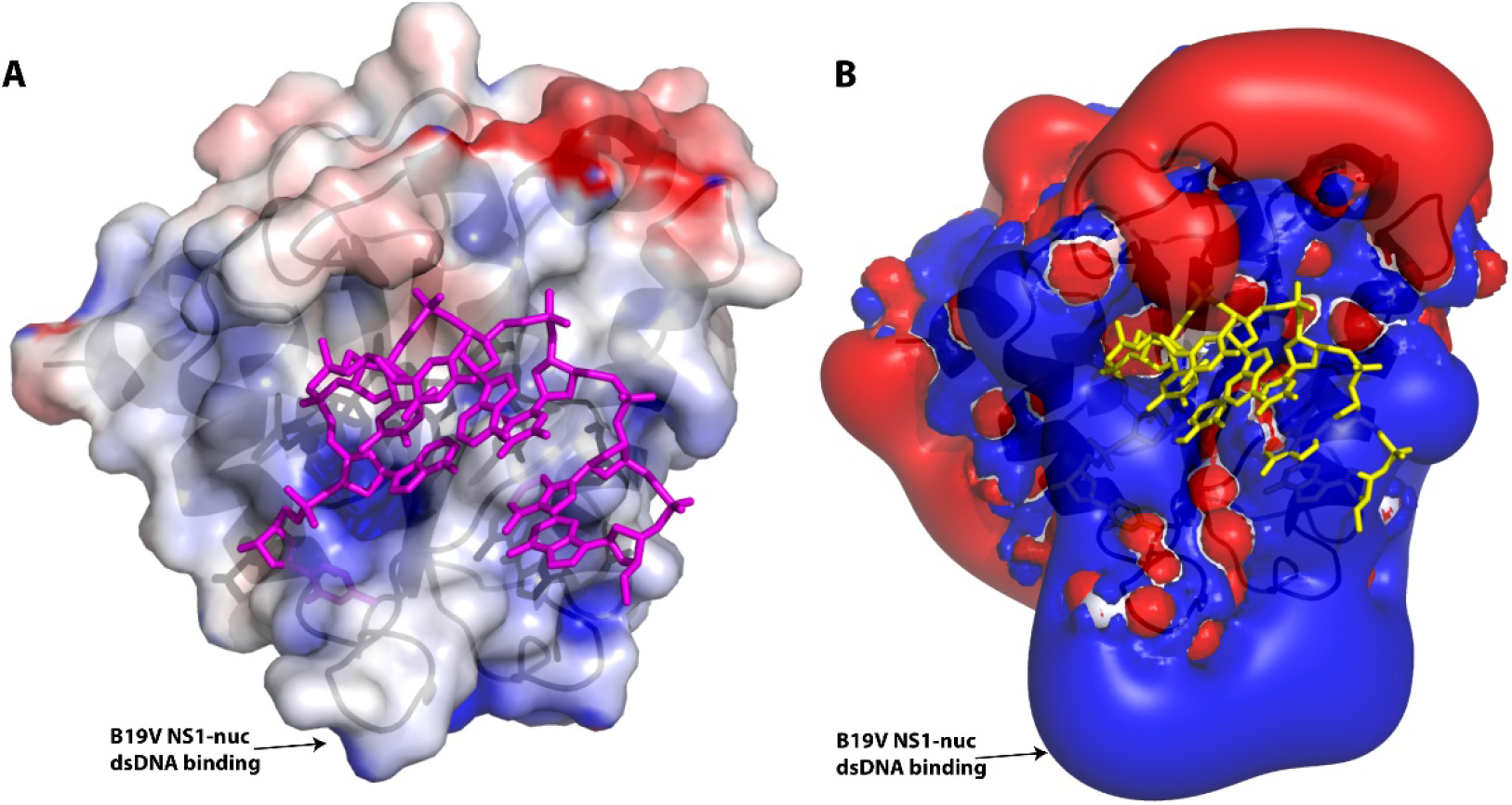
Electrostatic surface and potential map of NS1-nuc and predicted ssDNA binding site. **A.** Electrostatic potential of Form I NS1-nuc calculated using APBS^*39*^ in Pymol mapped onto the surface of Form I NS1-nuc. ssDNA from PDB accession file 6WE1^*45*^ after superposition of NS1-nuc and WDV Rep-nuc shown in magenta to mark the predicted ssDNA binding cleft. **B.** Electrostatic potential field calculated using APBS^*39*^ in Pymol of Form I NS1-nuc, with ssDNA from WDV Rep-nuc/ssDNA of PDB accession code 6WE1^*45*^ shown in yellow.

**Figure S8.**
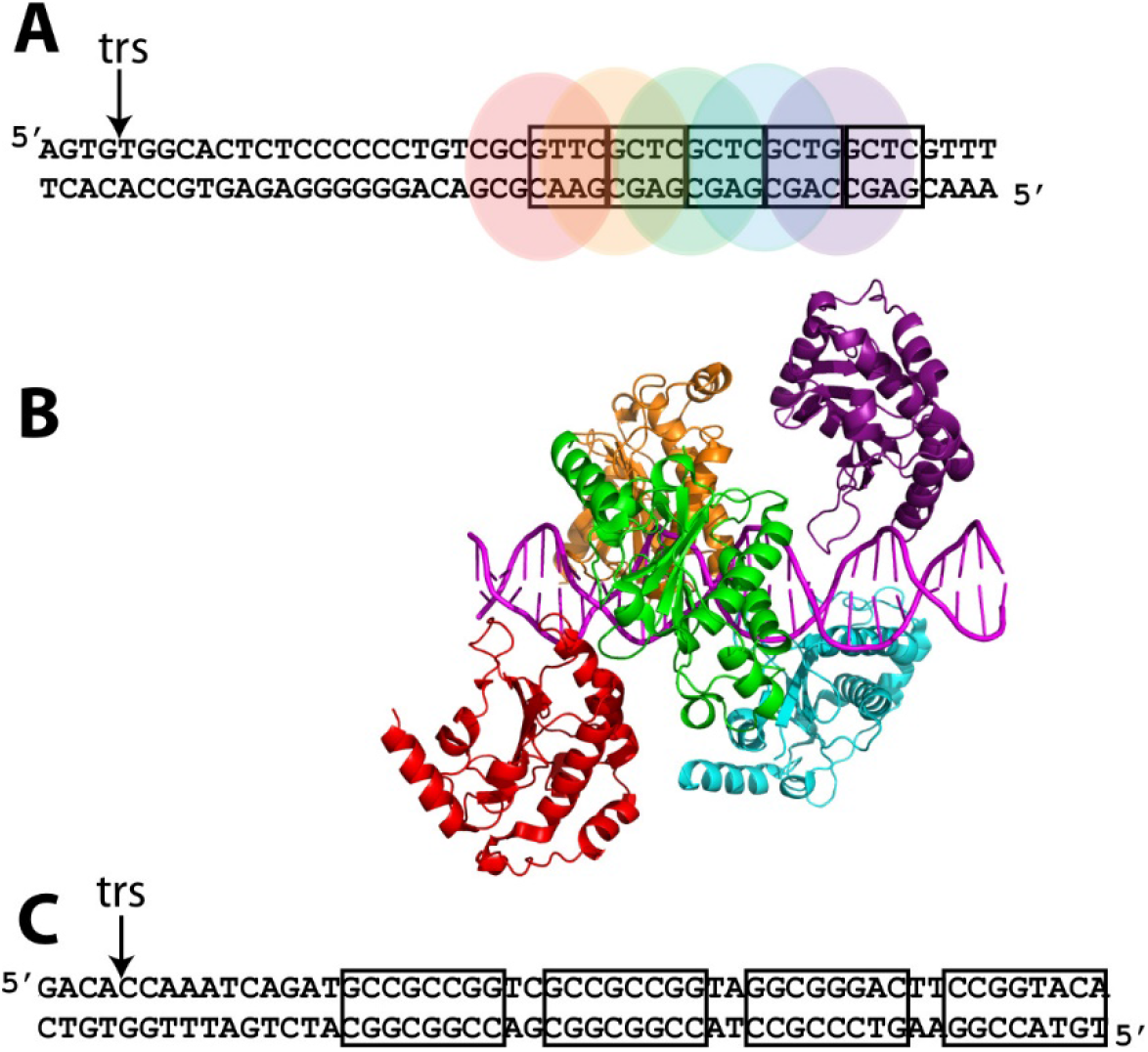
Interactions with dsDNA containing the viral origin of replication. **A.** AAV viral origin of replication sequence with RBE (boxed bp), trs (arrow, *a.k.a*. nicking site), and positions of each AAV Rep-nuc (from PDB accession code 1RZ9^*57*^). The position of each copy of AAV Rep-nuc is colored differently and corresponds to AAV Rep-nuc domain colors from the crystal structure shown in panel B. **B.** Ribbon drawing of five copies of the AAV Rep-nuc bound to RBE sequences in the AAV origin of replication (from PDB accession code 1RZ9^*57*^). DNA is shown in cartoon form and colored in magenta. **C.** B19V origin of replication sequences with NSBE (boxed) and nicking site/trs shown with arrow.

**Figure S9.**
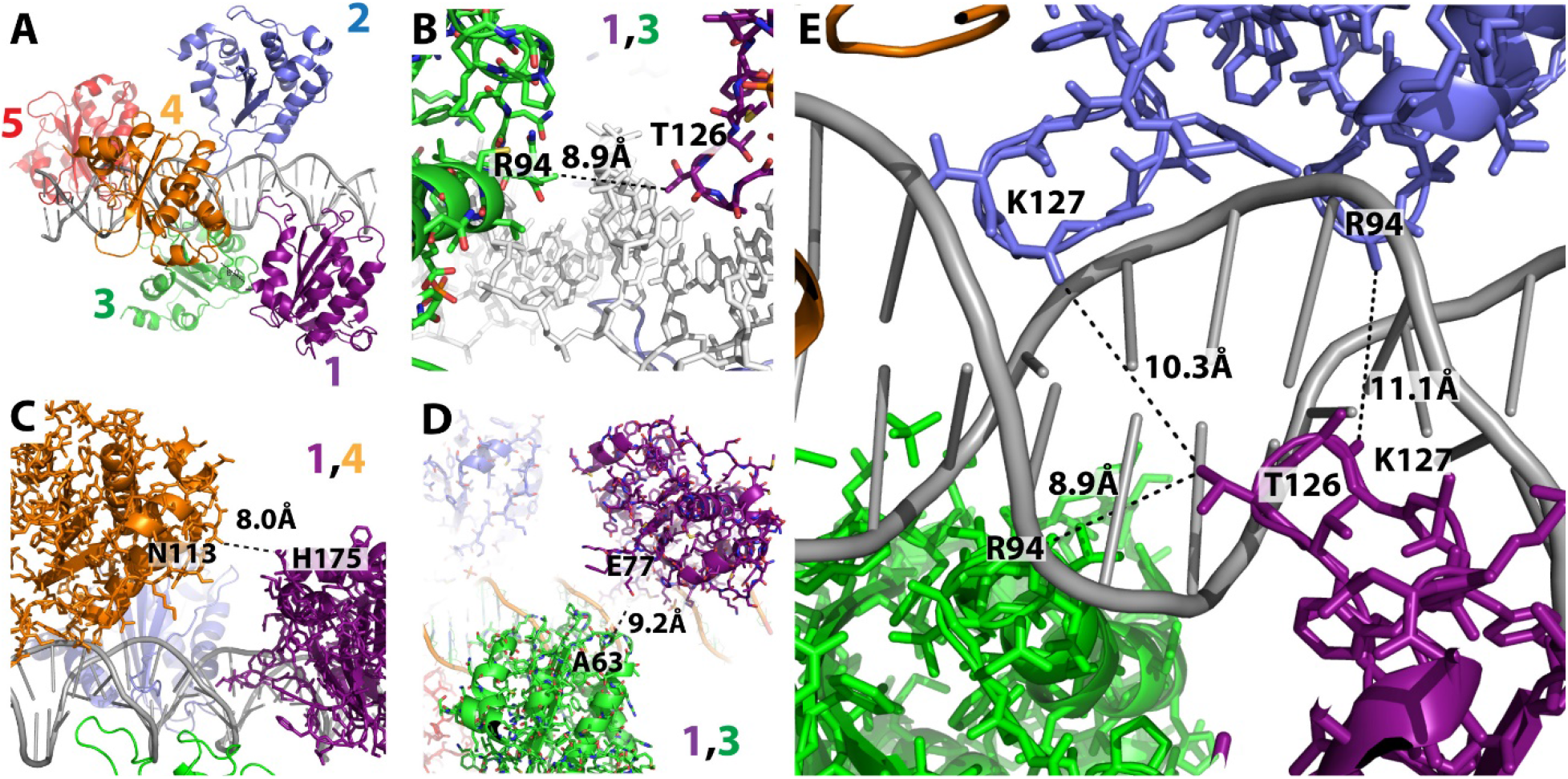
Closest approach of NS1-nuc molecules in dsDNA binding model based on AAV Rep-nuc/dsDNA crystal structure. **A.** Overview of NS1-nuc/dsDNA model based on PDB accession code 1RZ9^*57*^ containing five copies of the AAV Rep-nuc bound to RBE sequence dsDNA. DNA shown in grey cartoon form. Superpositions were created using all atoms of Form I NS1-nuc and each copy of AAV Rep-nuc. Each NS1-nuc domain binds in both the major and minor grooves, and subsequent copies follow the right-handed DNA helical twist with approximately four copies per turn. Each NS1-nuc copy is shown in a unique color and numbered according to DNA binding from right to left. **B.** Closest approach between copies 1 (purple) and 3 (green) occurs through the bound DNA. Note that the side chain atoms of R94 are not present in the model. **C.** Closest approach of copies 1 (purple) and 4 (orange). Note that H175 is the C-terminal residue of the model, and the side chains are absent. **D.** Closest approach between copies 1 (purple) and 3 (green) without any intervening atoms. **E.** Close approach of copies 1 (purple), 2 (blue), and 3 (green) through bound DNA. Note that side chains of R94 and K127 are not present in the model.

**Figure S10.**
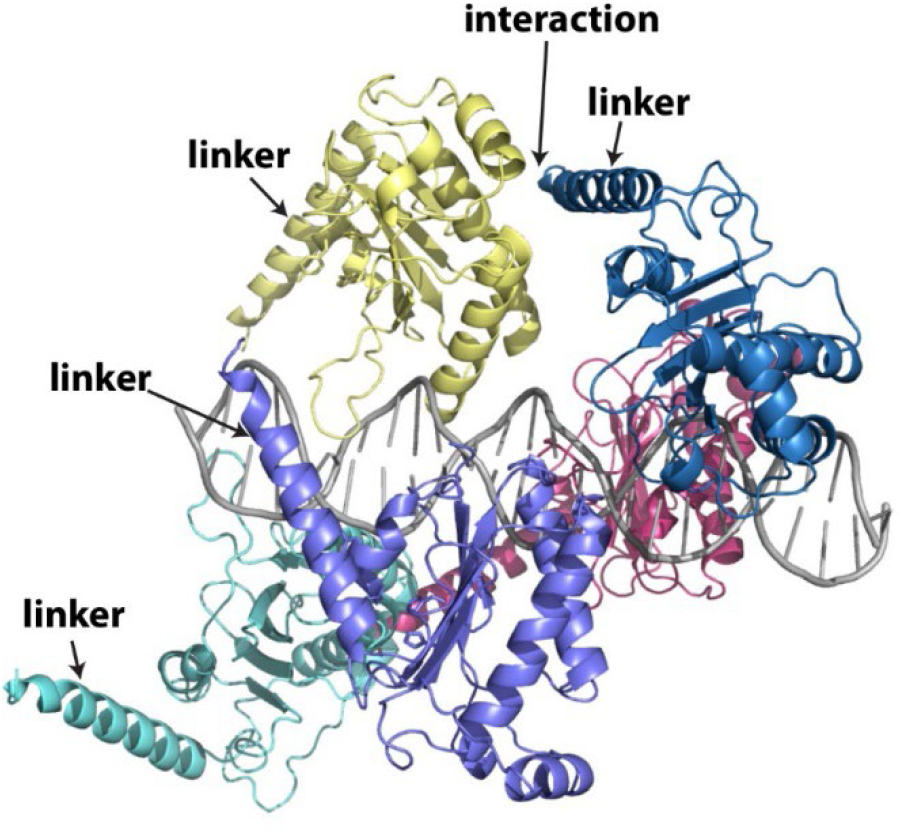
Model of AAV Rep-nuc containing additional C-terminal linker residues bound to dsDNA. The structure of AAV Rep-nuc containing residues of the linker region (identified with arrow) connecting the nuclease domain (PDB accession code 5DCX^*42*^) to the central helicase domain of AAV Rep superimposed onto AAV Rep-nuc domains found in PDB accession code 1RZ9^*57*^, which contains five copies bound to dsDNA. The linker approaches neighboring copies of AAV Rep-nuc such that protein-protein interactions may result.

**Figure S11.**
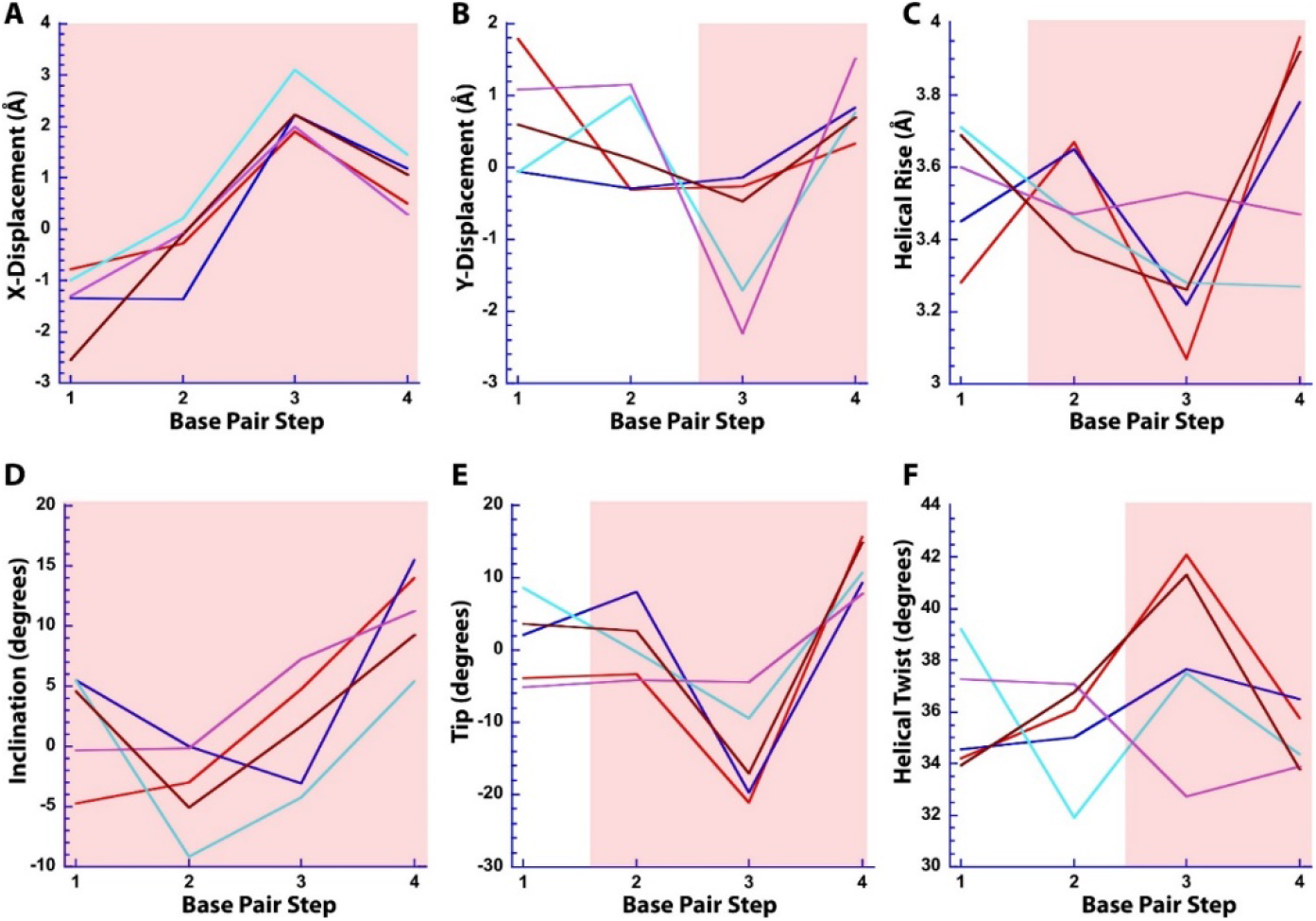
Helical parameters for dsDNA bound to AAV Rep-nuc. **A-F.** Helical parameters for the base pair steps of each RBE of the AAV origin of replication sequence (numbered left to right in Fig. S8A). RBE1: red, sequence: 5’GTTC, RBE2: blue, sequence: 5’GCTC, RBE3: cyan, sequence: 5’ GCTC, RBE4: purple, sequence: 5’GCTG, RBE5: brown, sequence: 5’GCTC. The fourth base pair step occurs between the last base pair of each RBE quartet and the neighboring base pair of the next RBE quartet. Red shaded area indicates helical deviation patterns shared by most RBEs.

